# Experience-induced remodeling of the hippocampal post-synaptic proteome and phosphoproteome

**DOI:** 10.1101/2021.10.26.465788

**Authors:** Seok Heo, Taewook Kang, Alexei M. Bygrave, Martin R. Larsen, Richard L. Huganir

## Abstract

The post synaptic density (PSD) of excitatory synapses contains a highly organized protein network with thousands of proteins and is key node in the regulation of synaptic plasticity. To gain new mechanistic insight into experience-induced changes in the PSD, we examined the global dynamics of the PSD proteome and phosphoproteome in mice following various treatments. Mice were trained using an inhibitory avoidance (IA) task and hippocampal PSD fractions were isolated for quantitative proteomic and phosphoproteomics analysis. We used a sequential enrichment strategy to explore the concurrent events of protein expression and phosphorylation in the hippocampal PSD following IA training (IA) or immediate shock (Shock). We identified more than 6,200 proteins and 3,000 phosphoproteins in the sequential strategy covering a total of 7,429 proteins. On the phosphoproteins we identified a total of 9,589 phosphosites. Strikingly, of the significantly IA-regulated proteins and phosphoproteins, a large fraction of the proteins displayed an overall decrease in phosphorylation level. Bioinformatic analysis of proteins and phosphoproteins that were regulated by IA were annotated for an involvement in regulation of glutamate receptor functionality, calcium signaling, and synaptic plasticity. We also identified synaptic kinases, phosphatases and their respective phosphosites regulated by IA training or immediate shock. Furthermore, we found that AMPA receptor surface expression was regulated by protein phosphatase, Mg^2+^/Mn^2+^ dependent 1H (Ppm1h). Together, these results unravel the dynamic remodeling of the PSD upon IA learning or immediate shock and serve as a resource for elucidating the synaptic proteome dynamics induced by experience-dependent plasticity.

**Highlights:**

- The proteome and phosphoproteome of mouse hippocampal PSD fractions were examined using quantitative phosphoproteomics and bioinformatics following inhibitory avoidance training or non-associative immediate shock.
- Approximately 6,200 proteins and 3,000 phosphoproteins were identified and quantified in the hippocampal PSD fractions.
- IA mediates widespread decreases in the abundance and phosphorylation of proteins in the hippocampal PSD fraction.
- Kinases, phosphatases and their phosphorylation status were dynamically and significantly regulated by IA and immediate shock.
- Functional validation shows that the protein phosphatase Ppm1h is linked to the regulation of synaptic plasticity *in vitro* and *in vivo*.

**In Brief:** Quantitative proteomics and phosphoproteomics combined with subcellular protein fractionation and bioinformatic analysis identifies a highly dynamic regulation of synaptic protein phosphorylation at the postsynaptic density following IA training and immediate shock.

## INTRODUCTION

The dynamic tuning of synaptic strength, through processes known as synaptic plasticity, is crucial for learning and memory (Huganir and Nicoll, 2013). Long-term potentiation (LTP) and long-term depression (LTD) are the two most studied forms of synaptic plasticity, arising from dynamic changes in neurons, including gene expression, protein trafficking and post-translational modifications (PTMs) (Costa-Mattioli et al., 2009; Ho et al., 2011). The post synaptic density (PSD) is an essential structure of excitatory synapses which is composed of both membrane and submembranous components. Biochemical and molecular biological studies have identified a number of proteins in the PSD, including neurotransmitter receptors, scaffold proteins, cytoskeleton proteins and signaling molecules, which together regulate synapse function, i.e., the communication between the pre- and post-synapses. Biochemical enrichment of PSD fractions and advances in proteomic approaches contribute to the expansion of our understanding of the synaptic proteins enriched in the PSD and their PTMs (Xu et al., 2021). However, the dynamics of PTM in the PSD upon various behavioral tasks in the brain, such as experience-dependent activity changes are poorly understood, and could potentially shed light on important signaling pathways for learning and memory processes in the brain.

LTP at the Schaffer collateral pathway between CA3 and CA1 pyramidal neurons in the hippocampus is the best characterized form of synaptic plasticity to date, both *in vitro* and *in vivo*. For example, LTP at CA3-CA1 synapses has been observed *in vivo* following inhibitory avoidance (IA) training in rats (Whitlock et al., 2006). The emotionally-motivated learning following single-trial IA training is hippocampus-dependent (Best and Orr, 1973) and robust and long lasting (Izquierdo et al., 1997). Memories formed by IA training are dependent on both protein synthesis and degradation. For example, studies have shown that inhibition of protein synthesis with protein synthesis inhibitors (e.g., anisomycin) infused in hippocampus or amygdala impaired consolidation, re-consolidation and extinction of IA-memories (Milekic et al., 2007; Taubenfeld et al., 2001; Vianna et al., 2001). In addition, proteasome-mediated protein degradation is also required for intact IA-memory (Fioravante and Byrne, 2011; Lopez-Salon et al., 2001). Protein phosphorylation/dephosphorylation of synaptic proteins also plays an important role in regulating the strength of synaptic connections (Diering and Huganir, 2018; Fingleton et al., 2021). The function of reversible protein phosphorylation mediated by kinases and phosphatases has been studied for decades and it is clear that phosphorylation is critically important for learning and memory (Coba, 2019; Woolfrey and Dell’Acqua, 2015). It is known that phosphorylation of different synaptic proteins is involved in different processes during memory formation (Lee, 2006; Woolfrey and Dell’Acqua, 2015). These data suggest that IA-learning and subsequent memory formation requires both synthesis and degradation of proteins, coupled with proper regulation of synaptic protein by PTMs such as phosphorylation.

Previous studies in mice have shown that IA induces changes in gene expression of c-Fos, Arc, homer1a, Na^+^/K^+^-ATPase subunits and glucose transporter type 1 (Tadi et al., 2015; Zhang et al., 2011). In rats (Cammarota et al., 1998; Whitlock et al., 2006) and mice (Chiu et al., 2017), IA training leads to recruitment of AMPA-type glutamate receptors (AMPARs) to the synaptosomal membrane fraction. In addition, GluA1 phosphorylation such as at the CaMKII site Ser831 is elevated following IA training (Cammarota et al., 1998; Whitlock et al., 2006). IA training also increased hippocampal CaMKII activity in the early phase of memory formation (Cammarota et al., 1998). These findings suggest that various synaptic proteins and their phosphorylation states are involved in IA-mediated learning and memory formation. However, our knowledge on synaptic proteins and their phosphorylation is limited to only a few synaptic proteins which are extensively studied.

Recent technological advances in mass spectrometry-based proteomics, including development of high-resolution mass spectrometry (HR-MS) instruments and tools for quantitative assessment of protein phosphorylation (Engholm-Keller and Larsen, 2016), alongside improvements in bioinformatics, enable unbiased characterization of proteins and their phosphorylation in brain with unprecedented depth (Bayes and Grant, 2009; Dieterich and Kreutz, 2016; Kempf et al., 2016; Kitchen et al., 2014; Palmisano et al., 2012). The power of proteomic approaches is being harnessed to identify how synaptic proteins and their phosphorylation change with learning when combined with behavioral testing and pharmacological manipulations (Borovok et al., 2016; Diering et al., 2017; Hong et al., 2013; Hosp and Mann, 2017; Kahne et al., 2016; Liu et al., 2018; McNair et al., 2006).

Here we used HR-MS combined with isobaric tags for relative and absolute quantification (iTRAQ) and TiO_2_-based phosphopeptide enrichment (Kang et al., 2019; Kang et al., 2018) to probe PSD-specific proteomic and phosphoproteomic remodeling following IA training. We identified a subset of significantly regulated PSD proteins and phosphoporoteins which showed decreased abundance 1 hour after IA training. Pathway analysis highlighted significantly enriched cellular functions related to normal synaptic plasticity, such as regulation of neurotransmitter receptor and ion transporter activity. Further analysis identified the involvement of distinct kinases and phosphatases (e.g., ppm1h), along with their phosphorylation sites, for the early phase of memory formation. This resource provides a novel perspective on IA- or immediate shock-associated hippocampal PSD proteome and phosphoproteome dynamics, revealing that large fractions of the synaptic proteins are differentially affected by different types of experiences.

## RESULTS

### Inhibitory avoidance training induces changes in glutamate receptors and their phosphorylation status

Initially, we sought to confirm that IA training induced a robust memory 1 hour after training, and that we could replicate changes in synaptic proteins that have been reported previously (Tadi et al., 2015; Whitlock et al., 2006). The IA training paradigm consisted of three sessions: 1) pre-testing habituation (5 minutes/day x 5 days), 2) training (IA, Walk, Shock groups), and 3) memory recall test. We used 4 experimental groups: IA trained animals (IA), a walk-through group that received no shock when crossing from the light to the dark (Walk), shock only (Shock) and naïve group (Naïve) animals (Figure 1A). One hour following training, IA memory was assessed by measuring the latency of mice to cross into the dark side of the chamber, after which mice were immediately euthanized and their hippocampi were harvested for PSD preparation (Figure 1B). As expected, during the training session, mice from both Walk and IA groups showed short latencies to cross to the dark chamber, indicating a preference for a dark environment (photophobia). In contrast, during the memory recall session, the IA group showed significantly longer latencies compared to the Walk group (Figure 1C). This demonstrates the robust one-trial learning induced by IA training.

**Figure 1.**
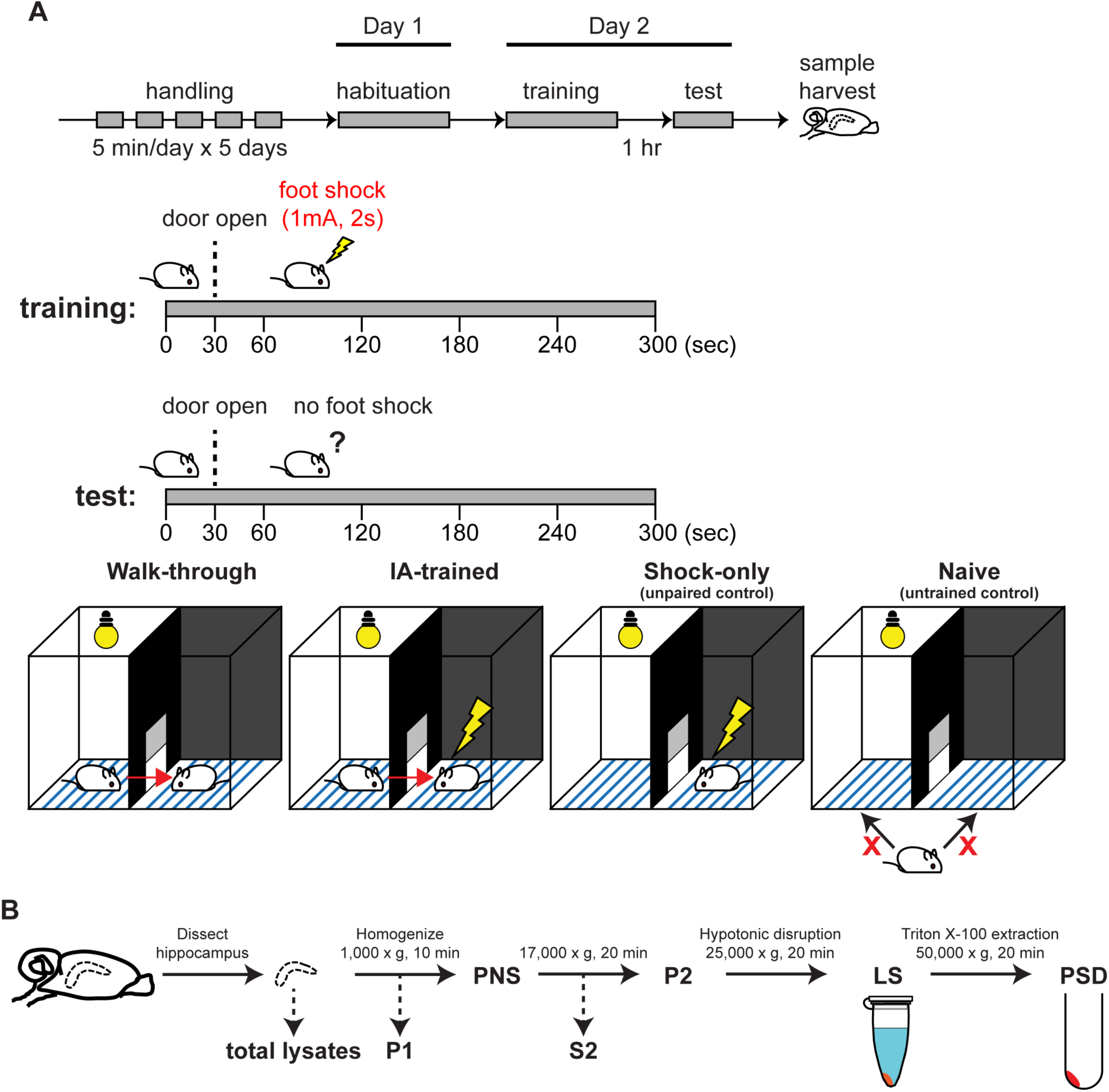
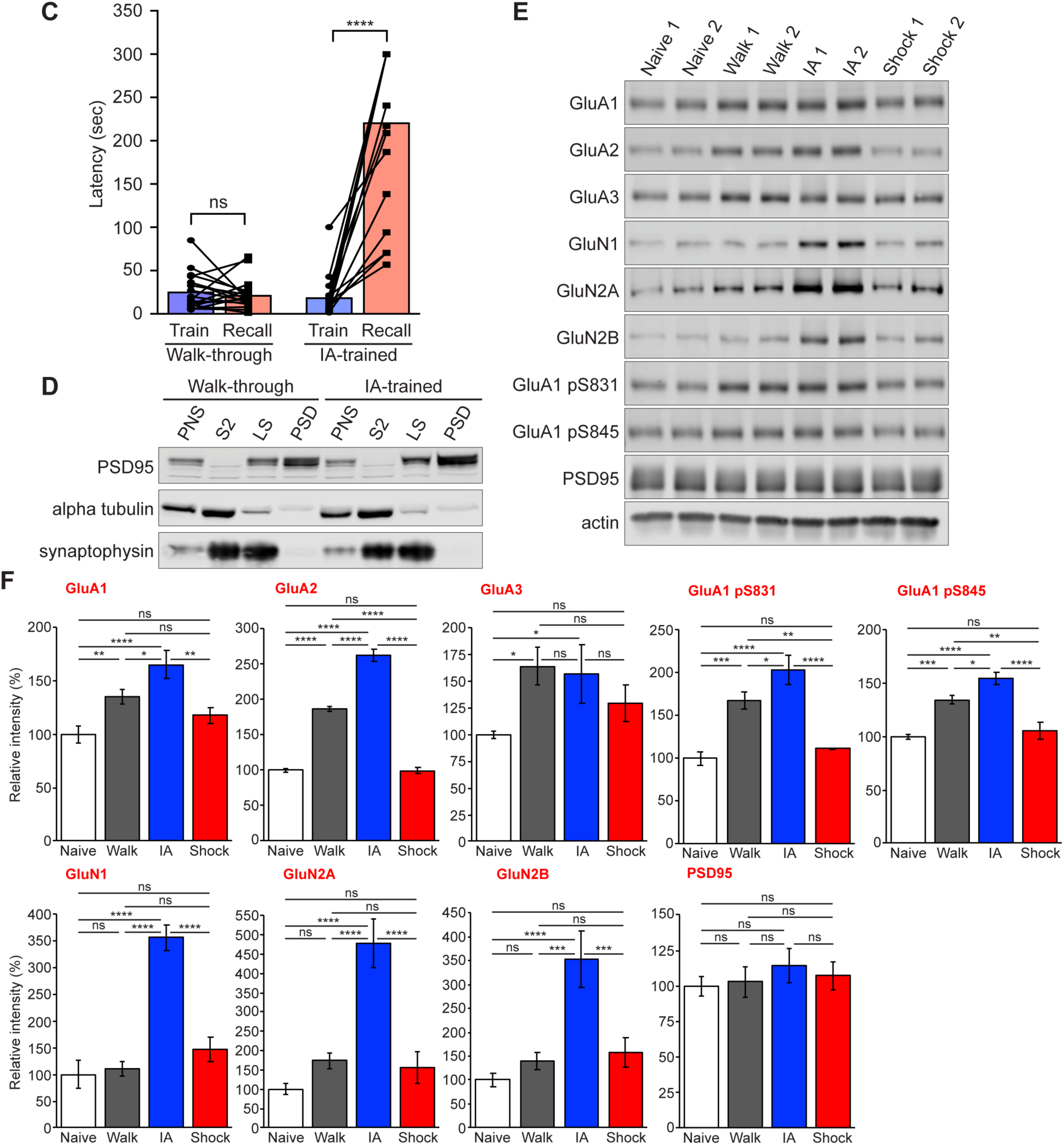
Experience-dependent dynamics of synaptic proteins. (A) Schematic of IA task. The behavioral test is composed of 3 sessions. Session #1 is handling, Session #2 is habituation, Session #3 is training and recall test followed by sample harvest (see STAR METHODS for details). (B) Schematic of subcellular fractionation for PSD preparation. (C) Behavioral results of IA training. The bar graph shows the latency for mice to cross to the dark side of the chamber, a measure of IA memory formation. Walk group (left) showed no significant changes in latency between train (blue) and recall test (red). IA group (right) showed significant increase of latency after training (2-way ANOVA, adjusted *p*-value < 0.0001). (D) Validation of the PSD fraction quality in representative mice from Walk (left) and IA (right) groups. Note the enrichment of PSD-95 and exclusion of alpha-tubulin in the PSD fraction. (E and F) Representative Western blots quantification of key synaptic proteins and their phosphorylation status in the PSD fraction. Western blot analysis to show changes of AMPAR, NMDAR and phospho- AMPAR. Error bars display ±SEM of 4 biological replicates.

Hippocampi harvested after the recall test were homogenized to prepare PSD fractions. Crude synaptosomes obtained from the post-nuclear supernatant (PNS) fraction were disrupted by hypotonic solution followed by PSD extraction using Triton X-100 (see STAR METHODS; Figure 1B). The quality of the PSD fraction was monitored by visualizing the enrichment of PSD95 and depletion of α-tubulin and synaptophysin in PSD fractions compared to other intermediate fractions (Figure 1D). Numerous studies have demonstrated that trafficking of different types of glutamate receptors contributes to LTP and other types of synaptic plasticity induced by learning (Bevilaqua et al., 2005; Lee et al., 2000; Lee et al., 1998; Lussier et al., 2015; Mitsushima et al., 2011; Park et al., 2014; Roth et al., 2017; Shipton and Paulsen, 2014; Zhang et al., 2015). We probed for changes in AMPA and NMDA receptors, and their phosphorylation status, in hippocampal PSD fractions from control (Naïve, Walk and Shock) and IA-trained mice. We found a significant increase of GluA1, GluA2 and GluA3 following IA training compared to the naïve control group. Subunits of NMDA receptors, GluN1, GluN2A, GluN2B, also increased following IA training. The well characterized phosphorylation sites of GluA1 at Ser831 (pS831) and Ser845 (pS845) increased compared to all control groups (Figure 1E and 1F). All AMPA and NMDA receptor subunits that we tested showed a robust increase in PSD following IA training.

Interestingly, GluA1, GluA2, GluA3, pS831 and pS845 of GluA1 also increased in Walk group compared to naïve and Shock group (Figure 1E and 1F). In contrast, no changes in the level of the synaptic scaffolding protein PSD-95 levels were detected. This validation indicates that IA training increases the targeting and phosphorylation of AMPA and NMDA receptor to the PSD, presumably underlying the expression of LTP *in vivo*.

### Quantitative analysis of the PSD proteome and phosphoproteome

To identify and characterize changes in PSD proteins and their phosphorylation status and potential signaling mechanisms mediated by IA training, we performed quantitative proteomics and phosphoproteomics followed by bioinformatic analysis in mice that underwent IA training (see STAR METHODS; Figure 2A). Proteins detected in more than two biological replicates were retained for subsequent analysis with various bioinformatic tools. From our master dataset, comprising of Naïve, Walk, IA and Shock groups, we successfully identified a total of 6,229 proteins and 3,038 phosphoproteins from PSD fractions, resulting in a total of 7,492 proteins identified with an overlap of 1,775 proteins (Figure 2B; Table S1). We next analyzed phosphoproteins identified and quantified from the PSD fractions. In the PSD fractions we identified a total of 8,954 unique phosphopeptides carrying 9,598 unique phosphosites on 3,038 phosphoproteins (Figure 2C; Table S1). Among these, we quantitatively compared 2,540 overlapping phosphopeptides (28.37%) carrying 2,621 phosphosites (27.31%) on 1,048 proteins (34.50%). Phosphopeptides that were detected at least twice in the triplicate experiments were considered for subsequent bioinformatics analysis.

**Figure 2.**
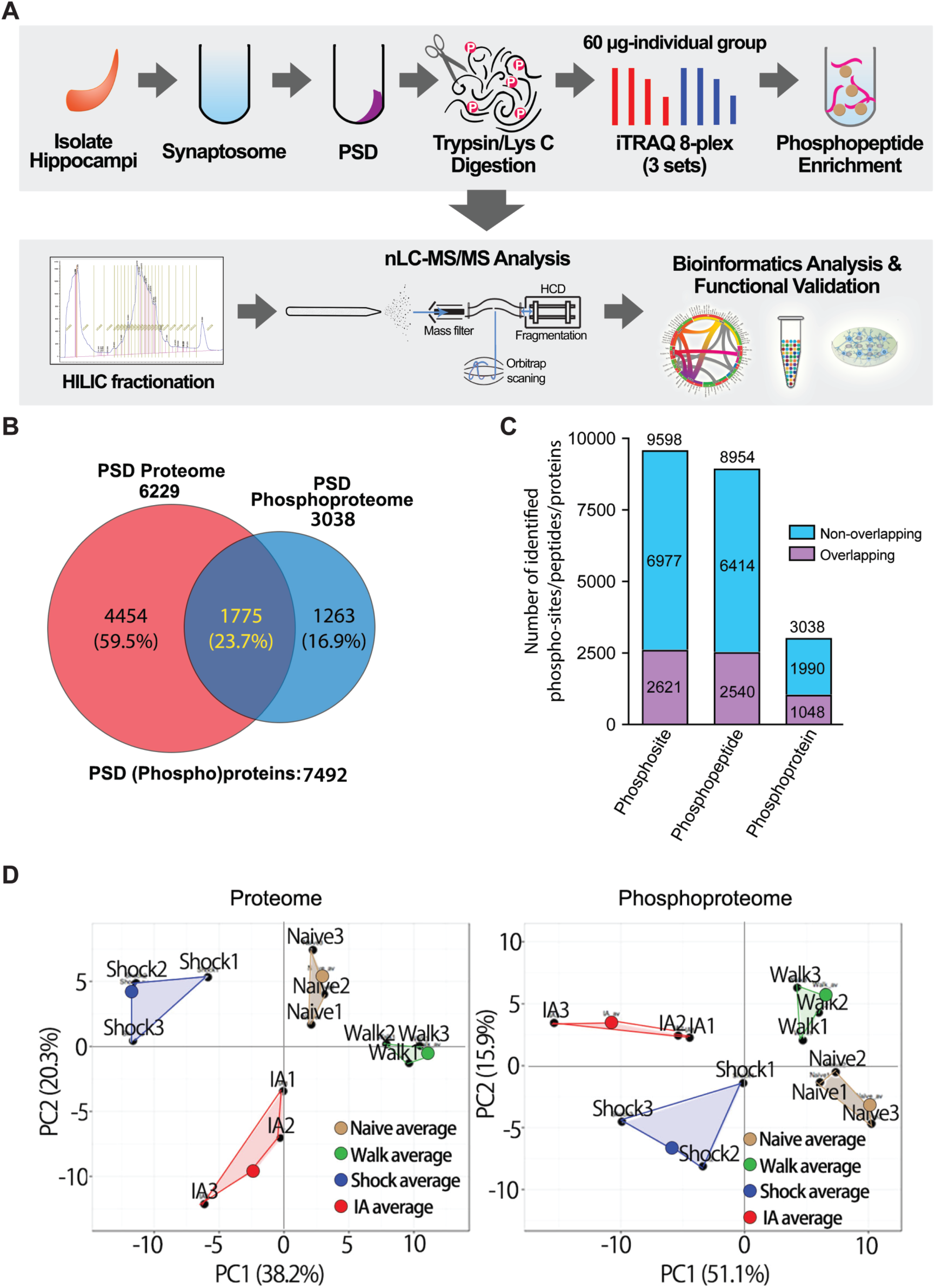
Identification and quantification of experience-dependent proteome and phosphoproteome dynamics in hippocampal PSD fractions. (A) Workflow of sample preparation and mass spectrometry (MS)-based phosphoproteomics analysis. PSD fraction was prepared from hippocampi dissected from individual mice from all 4 groups. Proteins were extracted and digested with trypsin/LysC to generate peptides for iTRAQ labelling. Multiplex labelled peptide mixture was subjected to phosphopeptide enrichment procedure using titanium dioxide (TiO_2_) beads. The flow-through (nonmodified peptides) and bound (phosphorylated peptides) fractions were desalted on R3 stage tip column and subsequently fractionated by hydrophilic interaction liquid chromatography (HILIC) fractionation. All fractions were analyzed using nLC-MS/MS. Acquired raw MS datasets were processed using MS-GF^+^ pipeline for protein identification and quantification followed by bioinformatics analysis and functional validation. (B) Venn diagram showing profile of mouse hippocampal PSD proteome and phosphoproteome identified and quantified from this study. (C) Bar chart showing the number of phosphosites, phosphopeptides and phosphoproteins identified from the hippocampal PSD fractions. Purple bars indicate overlapping phosphosites, phosphopeptides and phosphoproteins from biological replicates whereas cyan bars indicate non-overlapping ones. (D) Two-dimensional principal components analysis (PCA) comparing regulated nonmodified (left) and phosphorylated proteins (right) from IA (red), Walk (green), Shock (blue) and naïve (brown) groups based on component 1 and 2, which accounted for 38.2/20.3% for nonmodified proteins and 51.1/15.9% for phosphoproteins of the variability, respectively.

For an overall assessment of proteomic or phosphoproteomic similarities or differences of the four groups (naïve, Walk, Shock, and IA training), we used principal component analysis (PCA; see STAR METHODS). In a PCA from the PSD proteome, component 1 and 2, which account for 38.2% and 20.3% of total variability, respectively, clearly segregated each group into 4 distinct clusters (Figure 2D). Furthermore, the distance between the biological replicates is smaller than the separation between the groups. The PCA plot derived from the phosphoproteome data show similar results. Together with Western blot validation of various synaptic glutamate receptors in the PSD fractions (Figure 1E and 1F), these results further validate that the proteome and phosphoproteome changes in these groups were clearly segregated, supporting the modulation of proteins and phosphosites upon IA.

Based on this finding and the design of experimental groups reflecting exposure to the new environment (i.e., IA chamber), we used the Walk group as a control for comparative analysis between the IA and Shock groups.

### Experience dependent remodeling of the PSD proteome and phosphoproteome

We found that PSD levels of AMPA and NMDA receptors increased following IA training, and, albeit to a lesser extent, in the Walk group (Figure 1E and 1F). We hypothesized that the experience of exploring the inhibitory avoidance chamber, even in the absence of a shock (and associative emotional-learning), was enough to change neuronal activity and likely induce some changes in synaptic plasticity. Therefore, to better isolate learning-induced changes, we used the Walk group (with experience of the IA testing chamber) as an internal control in subsequent analyses, thereby comparing the abundance of proteins and phosphoproteins from IA and Shock groups to those from the Walk group.

Next, we hypothesized that IA training or immediate shock would comprise changes in the expression or phosphorylation levels of synaptic proteins that perform critical synaptic functions. To assess this, we analyzed the overall changes of the proteins and phosphopeptides in the PSD proteome in IA and Shock group compared to the Walk group. We observed that PSD proteins (n = 3,972) and phosphopeptides (n = 2,540) were regulated following IA training or immediate shock, and PSD phosphoproteome showed a broader distribution of changes than the proteome (Figure 3A). We next analyzed the trend of changes in expression and phosphorylation levels in the hippocampal PSD fractions following IA training or immediate shock. The degree of alterations of all quantified proteins showed insignificant changes in all 4 groups (Figure S1A). Interestingly, we observed that overall phosphorylation levels showed a decreasing trend in Shock and IA groups compared to the Walk group, with the IA group showing the greatest decrease in phosphorylation level (Figure S1B). The same analysis was performed using significantly regulated protein and phosphoproteins. We found that significantly regulated proteins from the IA group showed decreased expression levels compared to the other three groups (Figure 3B, upper panel). The decrease of protein levels is more obvious in the IA group, indicating that this decrease is somewhat task specific. Interestingly, PSD proteins in the IA groups showed the lowest levels of relative phosphorylation compared to Naïve, Walk and Shock groups (Figure 3B, lower panel). The overall phosphorylation decreases of approximately 45% was observed in the Shock group compared to the Naïve group. In comparison, in the IA group a phosphorylation decreases of approximately 60% was observed 1-hour post-training, while the phosphorylation decrease was only around 25% in the walk group.

**Figure 3.**
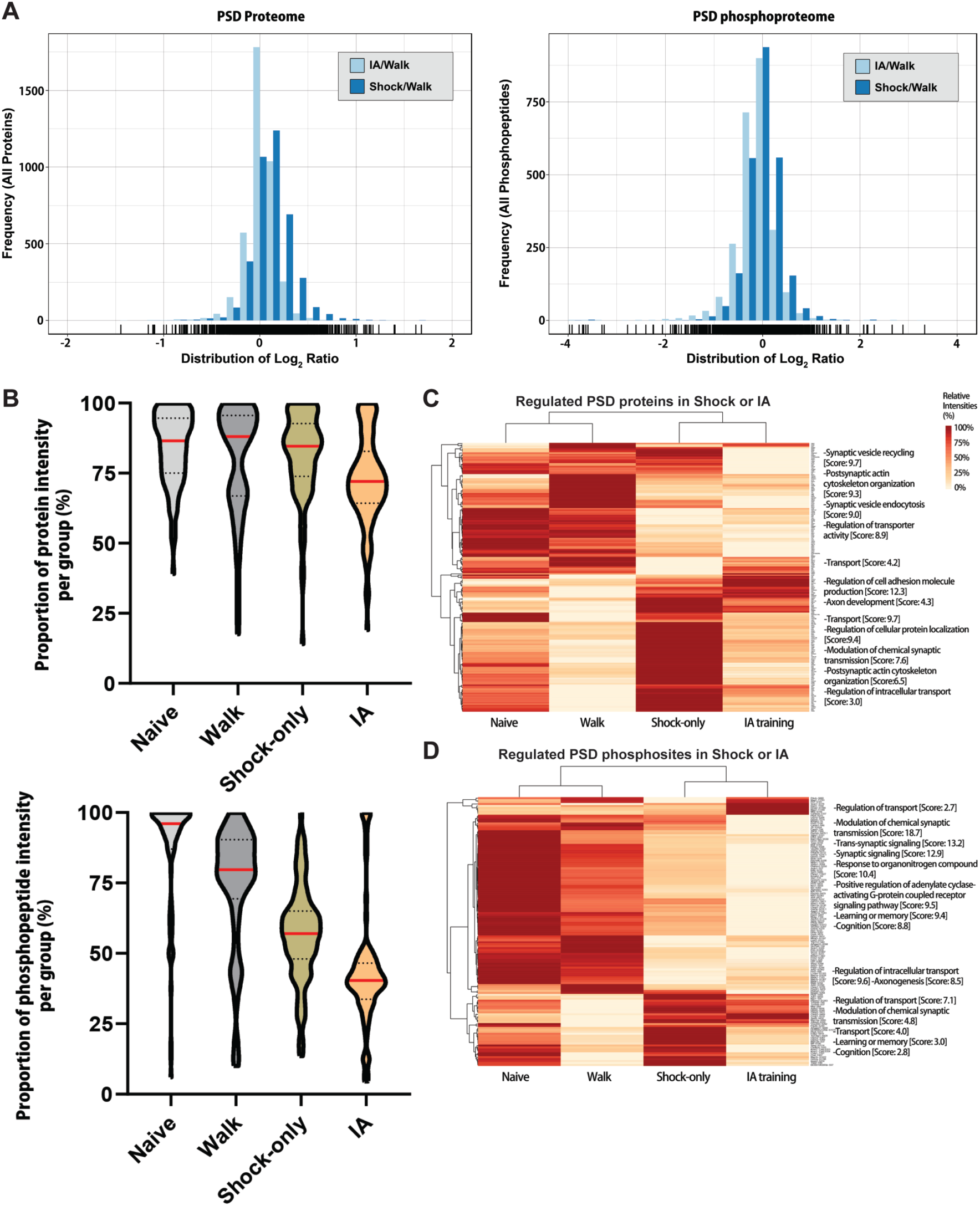
Dynamic changes of PSD proteins following IA-training and immediate shock. (A) Histogram of fold changes (Log_2_) showing frequency distribution of overall proteome and phosphoproteome regulation (including only overlapping proteins between biological replicates) from the IA or Shock group compared to the Walk group. In the comparison of IA group and Walk group, frequency of the nonmodified proteins and phosphopeptides are shown with light blue bars; frequency of the proteins and phosphopeptides from Shock group compared to the Walk group are shown with dark blue bars. (B) Violin plots showing the proportion of the significantly regulated proteins (left panel) or phosphopeptides (right panel) percentage based on normalized quantities in four groups following IA-training when compared to Walk group. Y-axis represents proportions of intensities of the regulated proteins (left) or altered phosphopeptides (right) intensities per group. Red lines inside of the violin plots represent the median of overall percentages per group. The width of the plot represents the density of proteins or phosphopeptides. Interquartile ranges are marked with dashed lines (Q3: upper quartile, Q1: lower quartile). (C) Heat map of the significantly regulated proteins (top) and phosphosites (bottom) showing the clustered patterns of experience-dependent regulation, for Naïve, Walk, IA and Shock groups (*n*=3 for each group) with hierarchical clustering (left side) and categories of biological functions obtained by IPA analysis (right side). The probability score based on *p*-value of biological function annotation are represented.

We visualized the relative percentages of normalized intensities of significantly regulated proteins and phosphoproteins in the PSD from each group with heatmaps combined with hierarchical clustering (Figure 3C). Proteins and phosphoproteins regulated by IA training or immediate shock were clustered based on their changing pattern across the groups followed by functional annotation analysis to reveal enriched biological processes. As shown is Figure 3B, we observed an overall reduction of protein and phosphorylation levels in the IA and Shock group. However, it should be noted that the function of individual phosphosites often is not known and therefore, the functional validation of selected phosphosites will be required to uncover the potential roles in synaptic plasticity or learning and memory. Analysis of proteins belonging to this cluster reveal significant enrichment of protein families involved in synaptic vesicle recycling, postsynaptic actin cytoskeleton organization, synaptic vesicle endocytosis and regulation of transporter activity (Figure 3C, upper panel). Analysis of proteins showing decreased phosphorylation levels in the IA and Shock group reveals significant enrichment of biological process terms, such as modulation of chemical synaptic transmission, trans-synaptic signaling, synaptic signaling, positive regulation of adenyl cyclase-activating GPCR signaling pathway, learning/memory, and cognition (Figure 3C, lower panel). Other clusters showing different patterns of regulation in Walk, IA and Shock groups share some biological processes, but also have distinct biological processes. Taken together, levels of proteins and their phosphosites are dynamically regulated by the different types of experience.

### Bioinformatic analysis of experience-dependent proteome and phosphoproteome dynamics

To analyze individual proteins and their phosphosites regulated by IA training or immediate shock, we grouped proteins and phosphoproteins based on their direction of change compared to the Walk group. We discovered 198 proteins that were significantly regulated following IA training or immediate shock (Figure 4A upper panel, IA↑/Shock↑: 89, IA↓/Shock↓: 80, IA↑/Shock↓: 4, IA↓/Shock↑: 25) and 138 phosphosites (Figure 4A lower panel, IA↑/Shock↑: 34, IA↓/Shock↓: 92, IA↑/Shock↓: 4, IA↓/Shock↑: 8). Among these regulated proteins, we identified 61 proteins and 57 phosphosites distinctively regulated by IA training. Figure 4A depicts all proteins and phosphosites analyzed, and each quadrant in the graphs shows four different categories of changes in the expression or phosphorylation levels of individual proteins. The upper right and lower left quadrants depict proteins and phosphoproteins that exhibit same directional regulation: both IA training and immediate shock result in either an increase (upper right quadrant, 89 proteins and 34 phosphosites) or a reduction (lower left quadrant, 80 proteins and 92 phosphosites) in the expression or phosphorylation levels (Figure 4A; Table S1) when compared to the Walk group. The remaining categories are represented by proteins and phosphoproteins regulated in a bi-directional manner. Enhanced protein or phosphorylation levels for these proteins are associated with one form of experience (either IA training or immediate shock) while reduced protein or phosphorylation levels are associated with the other form of experience. Proteins and phosphoproteins categorized in lower right quadrant (4 proteins and 85 phosphosites) were enhanced by IA training but reduced by immediate shock while those in upper left quadrant (25 proteins and 8 phosphosites) were reduced by IA training but enhanced by immediate shock (Figure 4A; Table S1). Taken together, we found dynamic remodeling of the PSD proteome and phosphoproteome following IA training or immediate shock compared to the walk-through control. To isolate changes unique to IA training or immediate shock, we looked for proteins and phosphoproteins that were regulated distinctively following IA training but not immediate shock or vice versa. Proteins and phosphosites regulated by single factor exclusively (IA training or immediate shock) are represented as spots close to x- (IA-unique) or y-axis (Shock-unique) on the scatter plot shown in Figure 4A (i.e., clustered within a ±5% window of the x- or y-axis). GO analysis of proteins and phosphoproteins regulated uniquely by IA training indicates a significant enrichment for proteins involved largely in the regulation of synaptic functions including regulation of postsynaptic neurotransmitter receptor and ion transmembrane transporter activity, regulation of adenylate cyclase-activating adrenergic receptor signaling, cellular Ca^2+^ homeostasis and modulation of chemical synaptic transmission (Figure 4B, left panel; Table S2). In the case of proteins and phosphoproteins uniquely regulated by immediate shock, GO analysis of this group indicates significant enrichment of cellular functions involved in positive regulation of LTD, regulation of Golgi organization, vesicle docking involved in exocytosis and negative regulation of microtubule depolymerization (Figure 4B, right panel; Table S2). Taken together, both IA training and immediate shock seem to engage some overlapping cellular functions, but there are also proteins and phosphoproteins that are regulated uniquely by IA training or immediate shock, which show distinct cellular functions in the GO analysis.

**Figure 4.**
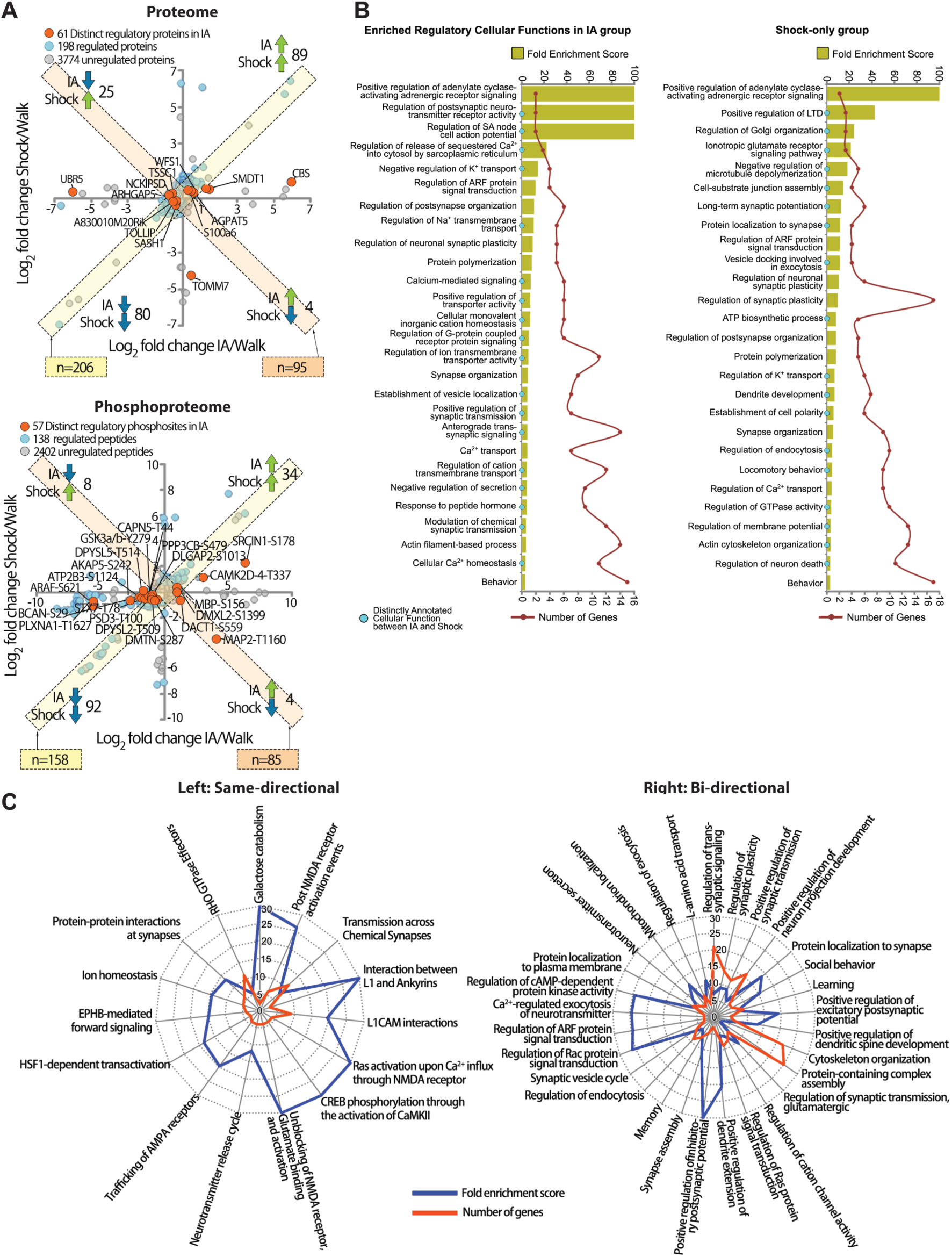
Bioinformatics analysis of PSD proteins and phosphoproteins regulated by IA-training and immediate shock. (A) Scatter plot showing all proteins (left panel) and phosphopeptides carrying relevant phosphosites in IA (x-axis) or Shock (y-axis) group compared to Walk group. Every analyzed protein (left, n = 4,033) and phosphopeptides (right, n = 2,597). Proteins and phosphosites that were significantly regulated by IA-training or immediate shock are indicated in light blue. Proteins and phosphosites that were significantly regulated by IA are indicated in orange. Other proteins and phosphosites that were not regulated are displayed in light gray. Proteins and phosphopeptides showing same- (IA↑/Shock↑ or IA↓/Shock↓) or bi- directional (IA↑/Shock↓ or IA↓/Shock↑) regulations are indicated in diagonal boxes with either light yellow or light orange color, respectively (CV ±5%). (B) Gene ontology (GO) enrichment analysis of regulated proteins and phosphoproteins in IA (left panel) or Shock (right panel) groups showing the fold-enrichment score (upper x-axis, gold bars) as well as the number of genes (lower x-axis, red line) for the indicated groups. PANTHER Gene Ontology analysis was performed to show enriched cellular functions (FDR < 0.05). Distinctly annotated cellular functions are marked with cyan dot on the left y-axis. (C) Radar plot showing signaling pathways on Reactome Pathway database (FDR < 0.05) that were affected by same- (left panel) or bi-directionally (right panel) regulated PSD proteins and phosphoproteins in IA and Shock groups. Fold enrichment of each pathway is displayed with blue lines, and the number of genes from each pathway is displayed with red lines.

Next, we conducted reactome pathway analysis (Jassal et al., 2020) to unveil high-order signaling pathways shared by the list of same- or bi-directionally regulated proteins and phosphoproteins following IA training and immediate shock. Proteins and phosphoproteins that were regulated with the same directionality in the IA and Shock groups revealed 15 significantly enriched pathways including Ras activation upon NMDAR-mediated Ca^2+^ influx, CREB phosphorylation through CaMKII activation, AMPAR trafficking, protein-protein interaction at synapses, and activation of NMDA receptors (Figure 4C left panel; Table S3). Proteins and phosphoproteins regulated bi-directionally in IA and Shock group showed 29 enriched pathways including positive regulation of excitatory/inhibitory postsynaptic potentials, dendritic spine development, dendrite extension, regulation of cAMP-dependent protein kinase activity, regulation of ARF and Rac protein signal transduction, and Ca^2+^-regulated exocytosis of neurotransmitters (Figure 4C right panel; Table S3). Indeed, neurotransmitter receptor-related reactome pathways were significantly enriched in the same directionally regulated protein group while bi-directionally regulated proteins and phosphoproteins revealed a broader spectrum of pathways involved in synaptic functions. Taken together, proteins and phosphoproteins that showed same- or bi-directional regulation by IA training and immediate shock might be regarded as ‘common’ or ‘specific’ proteins which were involved in shared or unique signaling pathways involved in IA training- or immediate shock-mediated plasticity, respectively.

### Clustering and mapping of protein interaction network related to IA-learning

Next, we employed fuzzy c-means clustering analysis to characterize the most enriched clustering patterns among differentially regulated proteins and phosphoproteins following IA training or immediate shock. We found that 102 regulated proteins and phosphosites were clustered in the group which decreased in the IA group compared to the Walk group (Figure 5A). We next used a combined approach of STRING (https://string-db.org/), DAVID (https://david.ncifcrf.gov/), and KEGG (https://www.genome.jp/kegg/) pathway analyses on the largescale dataset to systematically reveal protein-protein interaction networks among significantly regulated proteins and phosphoproteins (Kang et al., 2018). We found that significantly regulated proteins and phosphosites from PSD fractions in this cluster were closely connected (Figure 5B). We also found that relationship and individual molecular interactions of regulated proteins and phosphoproteins enriched 19 biological processes, including chemical synaptic transmission, actin filament-based process, axon guidance, cognition, regulation of GTPase activity and calcium-mediated signaling, which are associated with synaptic functions (Figure 5C).

**Figure 5.**
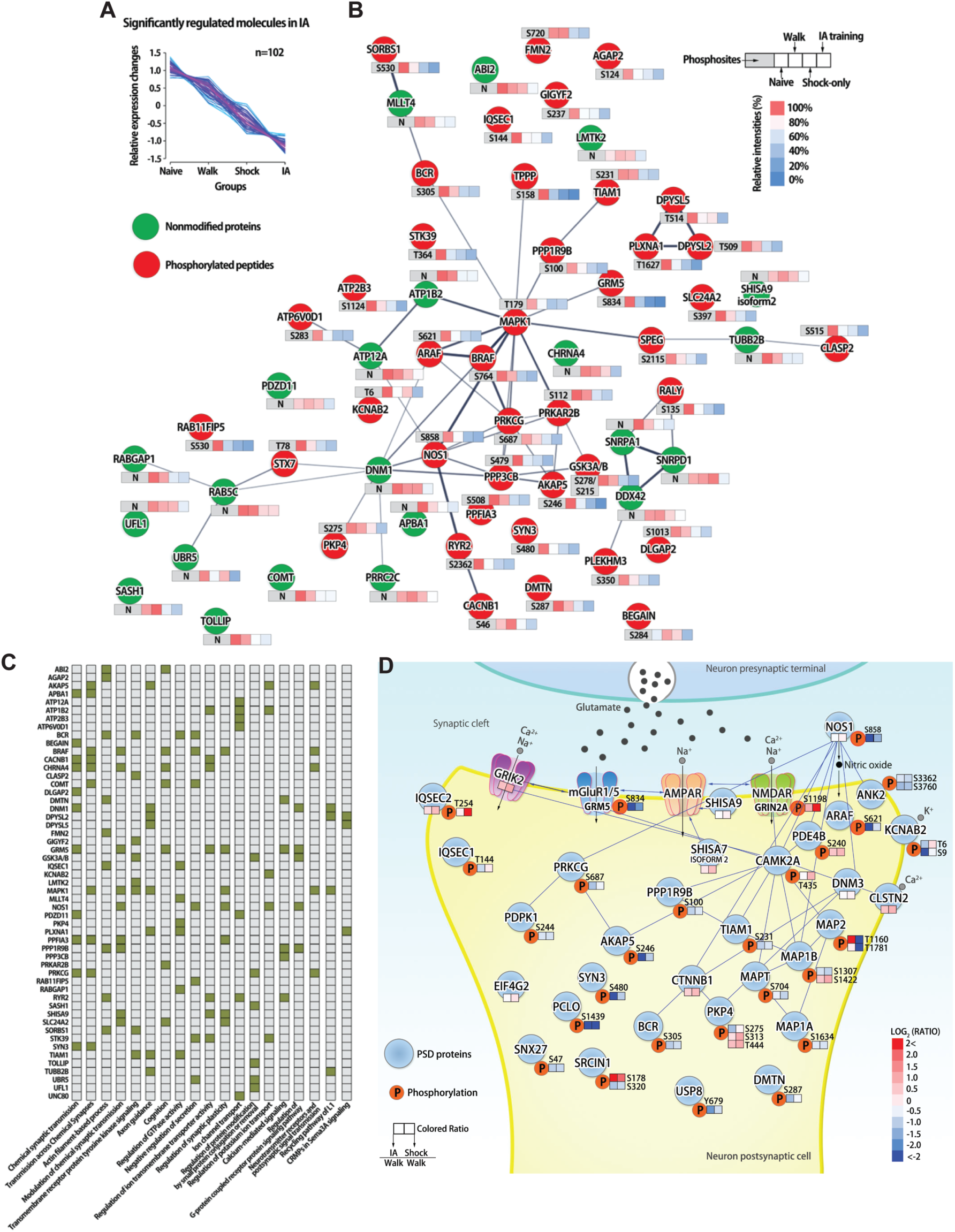
Protein-protein interaction network of regulated PSD proteins and phosphopeptides carrying relevant phosphosites following IA-training and immediate shock. (A) Most enabled clustering map of regulated proteins, phosphoproteins carrying relevant phosphosites that significantly decreased following IA-training (n = 102, adjusted p-value < 0.1 with CV ±30% and same directional regulation (negative) in all experiments) using unsupervised fuzzy c-means clustering analysis (membership > 0.6) of relative expression changes. Within this cluster, PSD proteins and phosphoproteins did not show significant changes following exposure to IA chamber (Walk). Immediate shock induced the trend of decrease in protein and phosphorylation level but it was not significant while IA-training induced a significant decreasing pattern of protein and phosphorylation levels. (B) Protein-protein interaction networks of the clustered proteins and phosphoproteins carrying relevant phosphosites showing a decreasing pattern following IA-training and immediate shock. The proteins and phosphoproteins from the most enabled cluster (significantly decreased in IA) were analyzed against STRING database to generate an interaction network map (confidence score > 0.7; high confidence). The quantitative information for each node (green: nonmodified proteins, red: phosphoproteins) is shown as relative concentration (0-100%) in a box format together with the information on designated phosphorylation sites. (C) Biological context of the interconnected PSD proteins and phosphoproteins with their relevant phosphorylation sites significantly decreased in IA. The gene symbols (left y-axis) that belong to enriched biological processes (x-axis) are indicated with colored square. (D) Schematic overview of the significantly regulated PSD proteins and phosphoproteins based on UniProt database following IA-training and immediate shock. The two squares represent the log_2_ ratio of the expression level change of designated proteins or their phosphorylation level in IA (left panel) or Shock (right panel) groups compared to the Walk group. Phosphoproteins are displayed with red-circled “P” and their phosphorylated site residue number.

We determined the changes that occurred in the post-synaptic compartment to find protein-protein interaction networks among PSD proteins from IA or Shock group classified from the postsynaptic density (GO:0014069). Within the post-synaptic compartment, we found several examples where IA training and immediate shock stimulation are associated with differential regulation of levels of proteins or phosphorylations in PSD proteins. For example, we found that neurotransmitter receptors and their phosphosites (e.g., GRIK2, S1198 on GRIN2A, S834 on GRM5) showed a pattern of increased protein abundance or phosphorylation in the Shock group (Shock/Walk) compared to the IA group (IA/Walk). We also found that phosphorylation levels on ion channel proteins (e.g., KCNAB2), kinases (e.g., CAMK2A, ARAF, PRKCG), chemical messenger producing enzyme (e.g., NOS1) and postsynaptic membrane proteins (e.g., CLSTN2) showed similar patterns of change (IA<Shock) in abundance or phosphorylation level. Other postsynaptic components that underwent significant downregulation in the IA group compared to the Shock group include IQSEC1/2, SHISA7/9 and AKAP5. Some proteins showed a significant increase in abundance or phosphorylation level in the IA and Shock groups. For example, levels of two phosphorylation sites of MAP2 (S1307, S1422) and SRCIN1 (S178) showed dramatic upregulation in the IA group, but showed significant downregulation in the Shock group. These proteins and phosphosites were interconnected with each other within the same protein interaction network map (Figure 5D). We also conducted signaling pathway analysis of regulated proteins and phosphoproteins to examine whether there is a functional relationship between IA training and immediate shock. We found that proteins and phosphoproteins regulated by IA training or immediate shock were significantly over-represented in signaling pathways including synaptic LTP/LTD, regulation of glutamate receptors, Rho family GTPase, synaptogenesis and actin cytoskeleton (Figure S2A: for regulated proteins, Figure S2B: for regulated phosphoproteins; Table S4). Within these protein interaction networks and signaling pathways, signaling molecules, such as protein kinases and phosphatases, were revealed as centers interconnected with other regulated proteins and phosphoproteins (Figure 5 and Figure S2). This result suggests that roles of these proteins and their phosphosites in signaling pathways are associated with experience-dependent remodeling of the synaptic proteome in the hippocampus after IA training or immediate shock.

### Protein interaction networks of significantly regulated proteins or phosphosites in hippocampus after both IA training and Shock

To investigate the changes of protein-protein interactions and their potential upstream proteins and their interacting proteins mediated by IA training or immediate shock, we performed fuzzy c-means clustering analysis using a combination of freely accessible search databases (e.g., IntAct, STRING, DAVID, Pubmed and UniProt) (Kang et al., 2020). First, we analyzed upstream interaction networks that were regulated in the same direction by IA training and immediate shock. We identified the eight most qualified upstream proteins showing changes in the same direction in both IA training and Shock groups (57 down-regulated and 43 up-regulated proteins/phosphoproteins) (Figure S3). Eight regulated upstream proteins including 14-3-3 protein families (YWHAZ, YWHAB, YWHAE), GRIN2B, PRKCE, KCNMA1, HSD3B4 and MACF1 were closely interconnected with each other. Interestingly, a large number of interacting proteins and phosphoproteins which potentially interact with 14-3-3 protein families were identified, and they were interconnected each other, which suggest that 14-3-3 may function as a signaling hub for experience-dependent proteome remodeling in the hippocampal PSD.

We determined the most qualified upstream proteins showing higher expression in the Shock group followed by manual confirmation of probable interacting proteins to reveal the major differences between the IA and Shock groups. The interactome map of immediate shock-induced changes in proteins and phosphoproteins showed six upstream proteins including YWHAB, RPGRIP1L, ANK2, GRIN1, GRIN2B and DLG4. These 6 upstream proteins regulated by immediate shock were grouped into three functional annotations; regulation of neuronal synaptic plasticity, regulation of Ca^2+^/K^+^ ion transport, negative regulation of GPCR protein signaling pathway (Figure S4). A series of DLG4 interacting proteins (proteins inside of the gray rim in Figure S4) interconnected with interacting proteins from YWHAB and GRIN1. Together with previous finding that 14-3-3 exert their activity by regulating NMDAR at postsynaptic sites (Qiao et al., 2014), this finding expands the relationship of NMDAR functionality with 14-3-3 at postsynaptic sites in the hippocampus as a response to immediate shock.

### Identification of regulated kinases and phosphatases

Protein phosphorylation is one of the most common PTMs, controlling important cellular processes through the action of kinases and phosphatases. Neuronal plasticity which mediates learning and memory also require different kinases and phosphatases that can reversibly phosphorylate and dephosphorylate specific sites on target proteins (Lee, 2006; Mansuy and Shenolikar, 2006; Woolfrey and Dell’Acqua, 2015). As shown in Figure 3 and 4, we observed dynamic changes in phosphorylation on several proteins, including various kinases and phosphatases, from the hippocampal PSD fraction. We performed sequence homology analysis (see STAR METHODS) of the significantly regulated kinases and phosphatases and their phosphosites to characterize the regulation pattern of individual enzymes and to identify novel kinases and phosphatases and their phosphosites associated with IA training or immediate shock. We found that 15 kinases (total number of kinases identified 2 out of 3 biological replicates: 241) were significantly regulated in the IA and Shock groups (Figure 6). These include 6 kinases, such as CAMK2A and 2D, PRKCG, PDPK1, MAPK1 and GSK3A/B, which are already known to be localized in the PSD and play crucial roles for synaptic plasticity (Cheng et al., 2011; Lisman et al., 2002; Lisman et al., 2012; Peineau et al., 2008; Saito and Shirai, 2002; Thomas and Huganir, 2004). We found that 9 kinases such as YES, LMTK, CSNK1E, ARAF, BRAF, STK39, AAK1, PRKCB isoform 2, and SPEG were newly identified in the PSD category using the QuickGO (https://www.ebi.ac.uk/QuickGO/term/GO:0014069). CAMK2 function as homomeric or heteromeric holoenzyme complexes, and each 12 subunits have different roles in synaptic plasticity (Coultrap and Bayer, 2012). We found that CAMK2A was up-regulated following immediate shock, but no obvious change was detected in IA group compared to Walk group. CAMK2D was down-regulated in both the IA and Shock group with the most dramatic decreased in the Shock group. The kinase phosphorylation pattern was differentially regulated compared to the total protein levels by different experiences. For example, phosphorylation of CAMK2A on T435 was up-regulated in the Shock group but slightly down-regulated in the IA group compared to the Walk group. However, CAMK2D phosphorylation on T337 was up-regulated, while the total CAMK2D protein level was down-regulated both in IA and Shock groups. Similarly, the total PRKCG protein level was up-regulated while phosphorylation on S687 was down-regulated in both IA and Shock groups. A series of kinases including PDPK1, GSK3A/B, AAK1, STK39, ARAF, CSNK1E showed higher expression in the Shock group compared to the IA group like CAMK2A. PRKCB isoform 2, MAPK1 and LMTK2 showed decreasing expression in both the IA and Shock groups (Figure 6, inside of the gray rim). Our results show that the expression patterns of different kinases are divergent depending on the experience.

**Figure 6.**
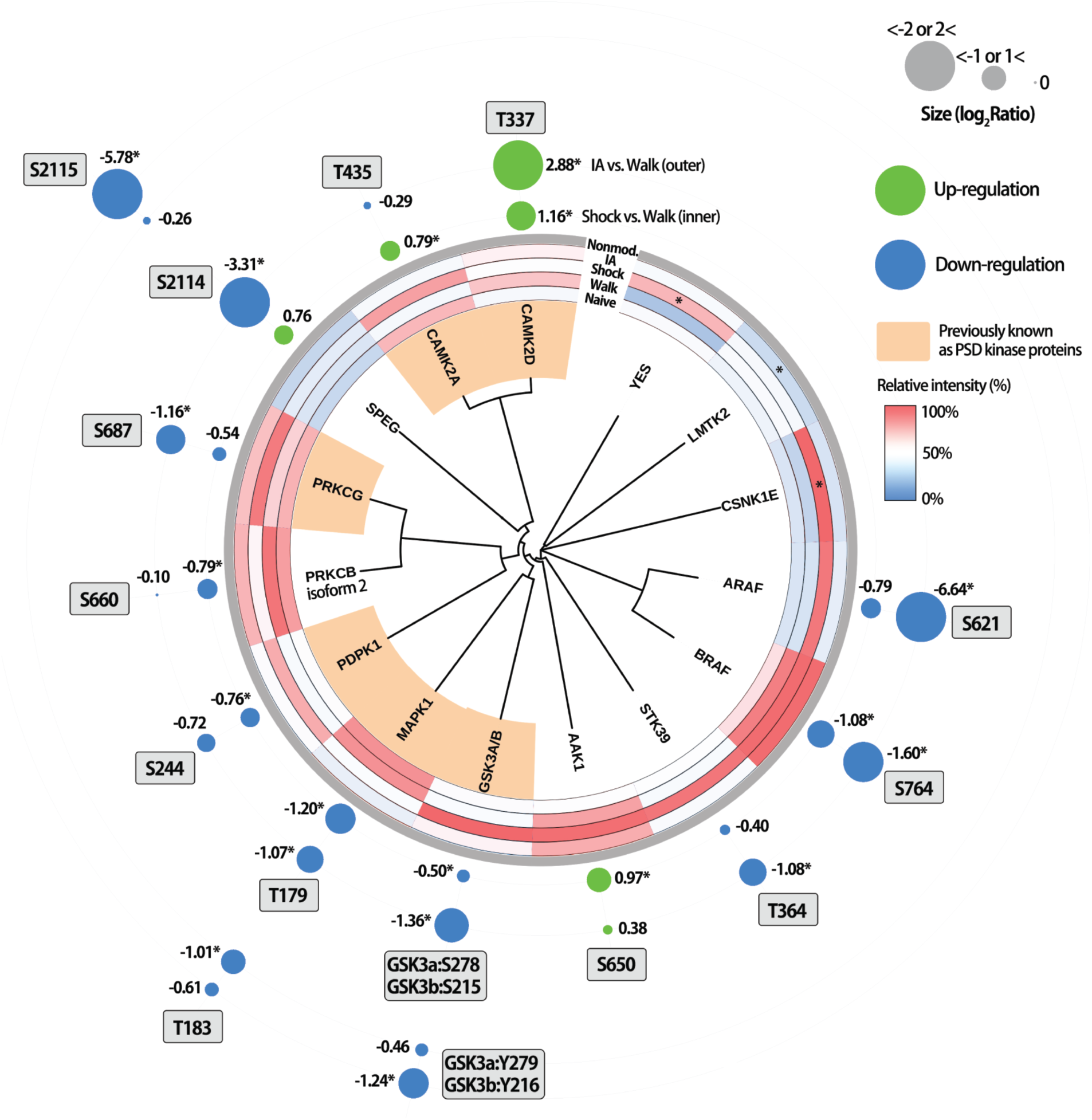
Profiling of regulated protein kinases. Kinome analysis of the significantly regulated kinases at the protein and phosphorylation levels in the PSD following IA-training or immediate shock with their homology of the kinase domains. Kinases that are known to exist in the PSD are labeled in orange. Relative intensities (%) of significantly regulated kinases following IA-training or immediate shock are shown in color-coded boxes (inside of the gray rim). The levels (log_2_ ratio) of phosphorylation of specific residues on individual kinases are shown as a circular index with different colors (green: up-regulated, blue: down-regulated) and size (outside of the gray rim). Outer and inner circles indicate Log_2_ ratio of given phosphosites in IA and Shock groups compared to Walk group, respectively.

The levels of autophosphorylation of regulated kinases also showed varied patterns. For example, phosphorylation at T435 of CAMK2A and S2114 of SPEG were found to be bi-directionally regulated by IA training or immediate shock (unfortunately, we did not identify the activity driven phosphosite in CAMK2A (T286) most likely due to the close tryptic cleavage sites). Phosphorylation of CAMK2D at T337 showed a significant increase in the Shock group and an even higher increase in the IA group. Phosphorylation of kinases which decreased both in the IA and Shock groups, such as SPEG (S2115), PRKCG (S687), GSK3A/3B (S278/S215, Y279/Y216), STK39 (T364), BRAF (S764), ARAF (S621), showed a greater decrease in phosphorylation following IA (Figure 6, outside of the gray rim). These results show that the expression levels of kinases and their phosphorylation levels are regulated differentially by experience.

### Functional validation of PSD proteins regulated by IA training or immediate shock

We observed a noticeable pattern of phosphorylation decrease in both the IA and Shock groups (Figure 3B). This finding led us to speculate about roles of protein phosphatases during IA training-mediated learning or in response to immediate shock. From our dataset, levels of protein phosphatases and their phosphorylation were found to be differentially regulated following IA training or immediate shock. Within the set of protein phosphatases regulated by IA training or immediate shock, we identified that phosphorylation of Ppm1h (protein phosphatase, Mg^2+^/Mn^2+^ dependent 1H), previously reported to be linked to synaptic plasticity (Bliim et al., 2019; Spiegel et al., 2014), was significantly down-regulated in both the IA training and Shock groups (Figure 7A). Unfortunately, we did not quantify the expression level of Ppm1h protein, although other isoforms of Mg^2+^/Mn^2+^-dependent protein phosphatase (e.g., Ppm1l, Ppm1b) were quantified (Figure 7A). Therefore, we additionally examined whether Ppm1h can be affected by different types of neuronal activity or regulate synaptic plasticity *in vitro* and *in vivo*. First, we electroporated *ppm1h* into cultured neurons and monitored the effect of Ppm1h overexpression on the levels of surface AMPA and NMDA receptors. Following Ppm1h overexpression, we observed a decrease of surface GluA1 GluA2, and phospho-GluA1 at S831 and S845, but no obvious changes in total expression level (Figure 7B). Interestingly, Ppm1h overexpression resulted in an increase of surface NMDAR subunits, GluN1 and GluN2A, but a decrease of surface GluN2B, indicating that Ppm1h likely regulates AMPARs and NMDARs differentially (Figure 7B). Next, we performed glycine-induced chemical LTP in cultured neurons and examined the abundance of Ppm1h (Figure 7C). The total Ppm1h level increased after 10 min glycine stimulation and was maintained during the 30 min chase period in the presence of Mg^2+^ (Figure 7D), suggesting that Ppm1h levels can also be regulated by chemical LTP.

**Figure 7.**
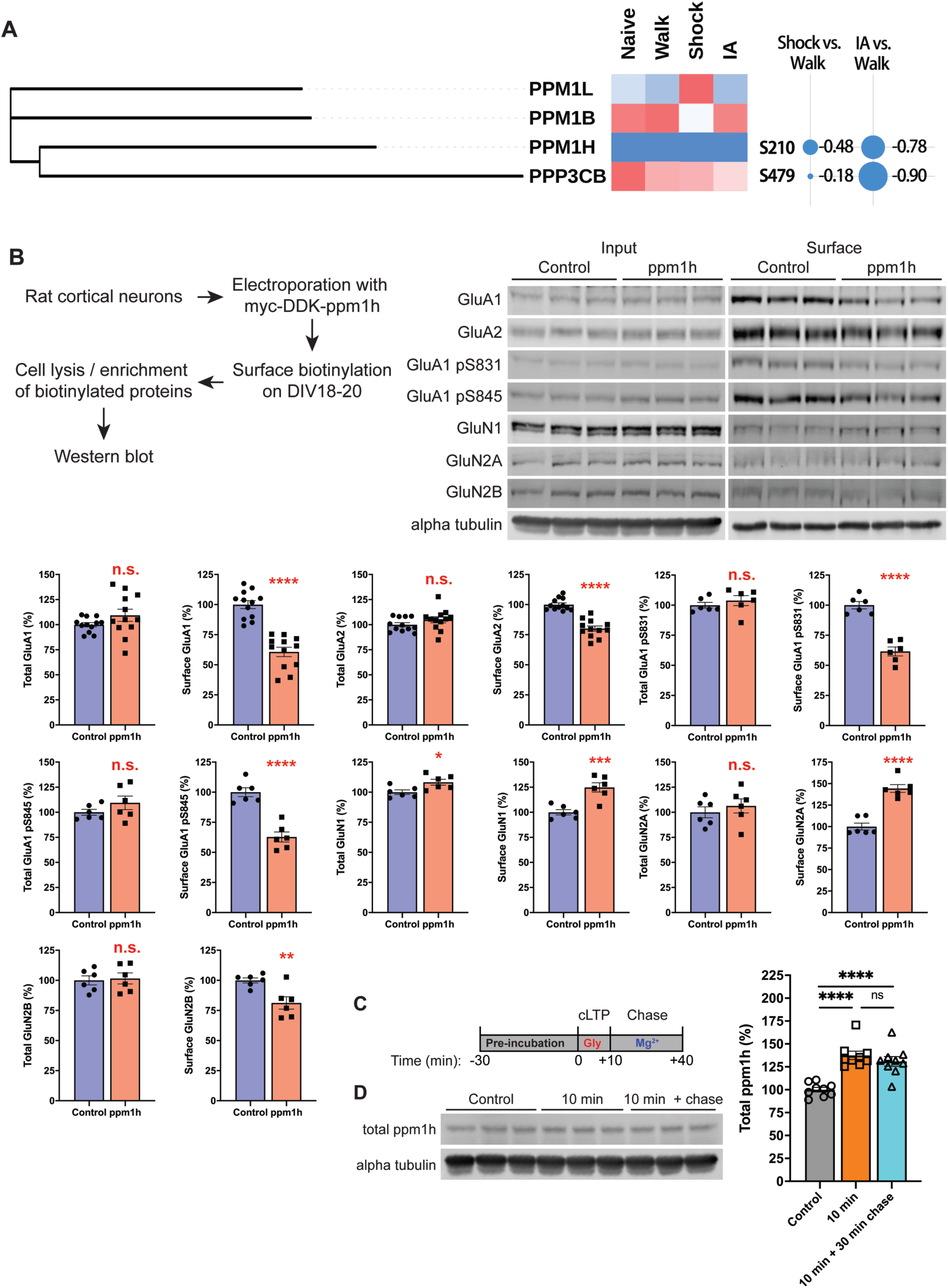
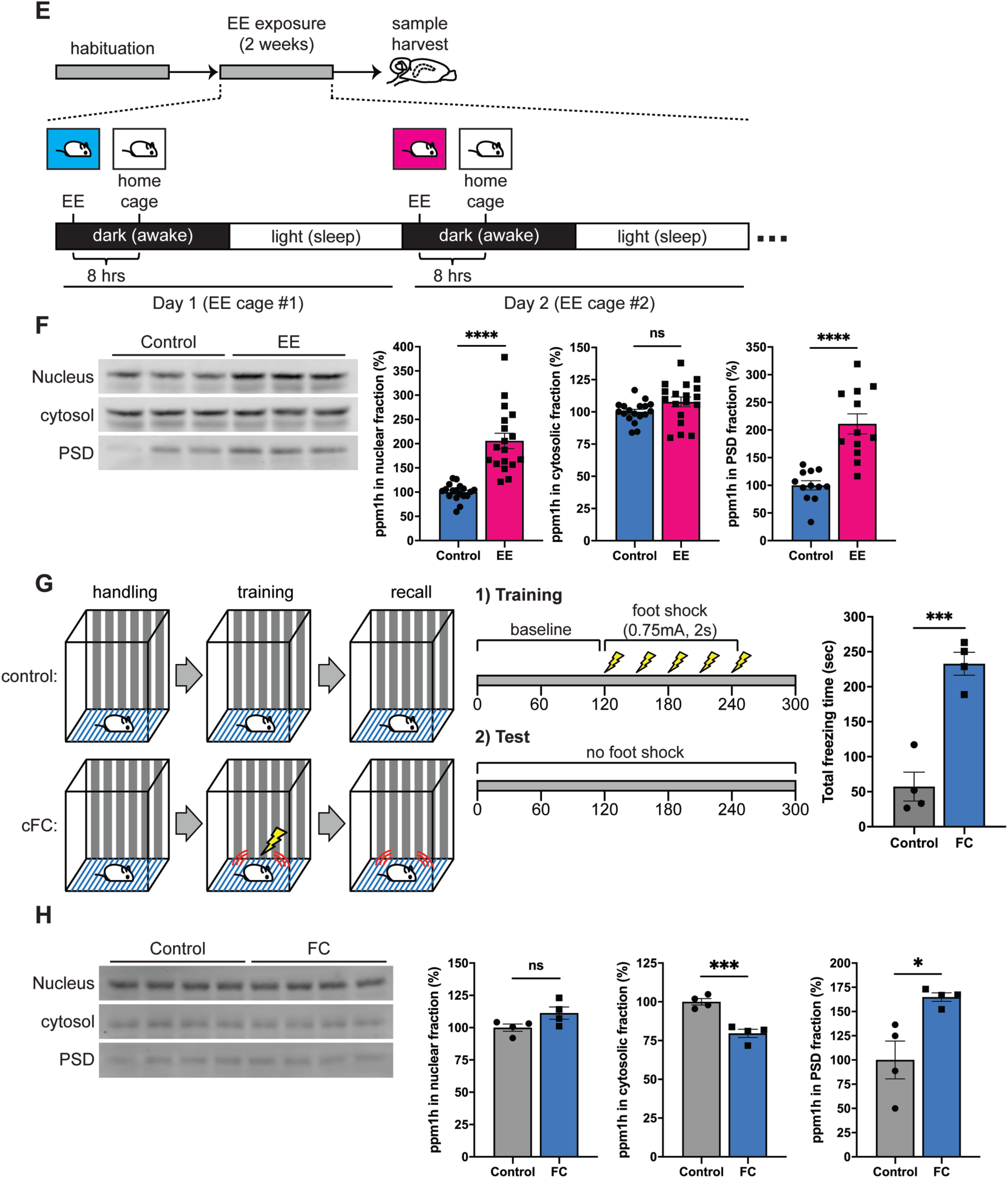
Functional validation of Ppm1h in synaptic plasticity. (A) Experience-dependent regulation of Mg^2+^/Mn^2+^-dependent protein phosphatases. Clustered isoforms of Ppm family showed unique regulations. Ppm1l had the highest level of expression Ppm1b the lowest expression following immediate shock. The phosphorylation levels on Ppm1h (S210) and Ppp3cb (S479) showed significant increase following IA-training. (B) Differential regulation of surface expression of AMPAR and NMDAR subunits by Ppm1h overexpression. When Ppm1h was overexpressed, cortical neurons exhibited increased surface expression of GluN1 and GluN2A. In contrast, surface expression of GluA1, A2, GluA1 pS831, pS845 and GluN2B showed a significant decreased following Ppm1h overexpression. (C and D) Regulation of Ppm1h during homeostatic scaling. (C) Scheme indicating experimental workflow for homeostatic scaling. Cultured cortical neurons (DIV11) were treated with 20 µM BIC or 1 µM TTX for 48 hrs to induce homeostatic down- or up-scaling, respectively. (D) Homeostatic scaling regulated expression levels of Ppm1h. TTX-mediated up-scaling exhibited slightly significant increase of Ppm1h level (1-way ANOVA, adjusted *p*-value = 0.025) while BIC-mediated down-scaling resulted in more significant increase of Ppm1h level (1-way ANOVA, adjusted *p*-value < 0.0001). (E and F) Regulation of Ppm1h during glycine-induced chemical LTP (GI-cLTP). (E) Scheme indicating experimental workflow for GI-cLTP. Cultured cortical neurons (DIV18 or older) were treated with 200 µM glycine for 10 min followed by chase with Mg^2+^- containing ACSF for 30 min. (F) GI-cLTP increased the expression level of Ppm1h levels after both 10 min stimulation and 30 min chase (1-way ANOVA, adjusted *p*-value for Control vs. 10 min and Control vs. 30 min chase < 0.0001). (G and H) Effect of enriched environment (EE) exposure to the level of Ppm1h in the mouse hippocampus. (E) Scheme indicating experimental workflow for exposure to EE. Mice had been tested for 2 weeks using 2-day EE exposure paradigm (see STAR METHODS), followed by isolation of subcellular fractions (nuclear, cytosolic and PSD fractions) from hippocampus. (H) Characterization of Ppm1h expression in different subcellular fractions following EE exposure. EE resulted in a significant increase of Ppm1h in the nucleus and PSD while cytosolic Ppm1h levels did not change. (I and J) Effect of contextual fear conditioning (cFC) to the level of Ppm1h in the mouse hippocampus. (I) Scheme indicating experimental workflow for cFC. Mice were trained via cFC task (see STAR METHODS), followed by isolation of subcellular fractions (nuclear, cytosolic and PSD fractions) from the hippocampus. Mice showed a significant increase of time spent freezing after cFC. (J) Characterization of Ppm1h expression in different subcellular fractions following cFC. Ppm1h significantly increased in PSD fractions while cytosolic or nuclear Ppm1h level showed a significant decrease or no change after cFC training, respectively.

Next, we investigated the regulation of Ppm1h following neuronal activity changes *in vivo*. First, we exposed mice to an enriched environment (EE) to induce brain-wide changes in neuronal activity and examined the levels of hippocampal Ppm1h from different subcellular fractions (Figure 7E; see STAR METHODS). We observed increased levels of Ppm1h from nuclear and PSD fractions while cytosolic fraction did not show any significant changes in the hippocampus after EE exposure (Figure 7F). Second, we similarly examined the levels of hippocampal Ppm1h from different subcellular fractions after contextual fear conditioning (cFC) to examine the effect of learning-specific neuronal activity changes (Figure 7G; see STAR METHODS). Interestingly, we observed a significant decrease of Ppm1h in the cytosol and an increase in PSD fractions while nuclear Ppm1h levels did not change (Figure 7H). Taken together with our quantitative proteomic results, these data indicate that levels of Ppm1h are differentially regulated by different types of neuronal activity and that manipulation of Ppm1h level can affect the trafficking of glutamate receptors to the surface of synapses, which may subsequently affect the regulation of synaptic plasticity *in vitro* and *in vivo*. This places de-phosphorylation as another important regulator of synaptic plasticity.

## DISCUSSION

In this study, we examined the dynamics of the proteome and phosphoproteome induced by IA training and immediate shock in mouse hippocampal PSD fractions by using high resolution mass spectrometry combined with a multiplexed quantitative proteomics approach. iTRAQ labeling combined with phosphopeptide enrichment using TiO_2_ enabled us to identify and quantify proteins and specific phosphosites in the PSD fractions from mice that were exposed to different experiences (i.e., walk-through, IA training and immediate shock). We identified and quantified over 6,200 PSD proteins and 3,000 phosphoproteins (including over 9,500 phosphosites). Alteration in the expression levels of PSD proteins and phosphosites were observed after IA training or immediate shock. These proteins and phosphoproteins were largely involved in neuronal functions, such as synaptic plasticity, regulation of neurotransmitter receptors, ion channels, and structural organization of synapses. One of the most interesting change was a significant decrease in protein and phosphorylation level followed by both IA training and immediate shock.

Recently, proteomic approaches are a critical tool to track the changes of proteome and post-translational modifications in the field of neuroscience (Li et al., 2017; Schanzenbacher et al., 2018; Thygesen et al., 2018; Thygesen et al., 2019). In this study, we analyzed the proteomic and phosphoproteomic remodeling that occurs in the hippocampus of mice following different types of experience to investigate molecular mechanisms underlying experience-dependent remodeling of synapses. We applied multiple control groups, including Naïve group, to distinguish the effect of learning on proteome remodeling by excluding the effect of other external stimuli, such as exposure to the new environment (Walk) or immediate aversive stimulation (Shock). We demonstrated that proteins and phosphosites are dynamically regulated following the robust learning induced by IA training. We also found that associative learning by IA training resulted in differential proteome and phosphosite remodeling compared to non-associative stimulation (immediate shock) as well as to the exposure to the new environment (walk-through). We provide comprehensive datasets highlighting experience-dependent remodeling of the hippocampal proteome *in vivo*. The derived lists of proteins and phosphoproteins from our quantitative proteomic analysis represent the patterns of proteome dynamics that may shed new light on the mechanisms underlying synaptic plasticity and learning and memory.

Synaptic plasticity is associated with the delivery of different types of glutamate receptors to the synapses (Huganir and Nicoll, 2013). In a previous study, phosphorylation of GluA1 at Ser831 increased, whereas phosphorylated GluA1 at Ser845 was not affected by IA training. Synaptic targeting of total GluA1 and GluA2 AMPA receptor subunits, but not NR1 NMDA receptor subunit, was enhanced after IA training (Whitlock et al., 2006). Our validation experiment showed a net increase of GluA1, GluA2, two well-characterized phosphosites of GluA1 (Ser831 and Ser845) and NMDAR subunits (GluN1, 2A, 2B) in PSD fraction after IA training (Figure 1E and F). Interestingly, we observed significant increase of AMPAR, NMDAR subunits and two well-characterized phosphorylation sites of GluA1 in the mouse hippocampal PSD fractions from Walk group compared to the Naïve group. Our approach is a short-term exposure to the training chamber (duration: maximum 5 minutes, number of exposures: maximum two times) followed by the immediate harvest of the hippocampus. We have previously shown that mice exposed to the enriched environment for 2 hrs exhibited increase of total and phosphorylated GluA1 at Ser831 and Ser845 in the mouse forebrain PSD fractions (Diering et al., 2016). Since the concept of enriched environment had been originally introduced by Donald Hebb (Brown and Milner, 2003), many studies showed that enriched environment has a considerable number of effects including gene expression, transcription and translation, throughout the brain (Kempermann, 2019). Although further studies need to be conducted to answer why AMPARs, NMDARs and phospho- GluA1 increased after short exposure to the training platform, our results suggest that appropriate cohorts which are exposed to the same behavioral apparatus without external stimulation, in this study the Walk group, should be set as a control group for memory assessment and biochemical validation.

Here, we employed IA task which is a commonly used behavioral task to investigate learning and memory processes (Cammarota et al., 1995; Tadi et al., 2015; Whitlock et al., 2006). This task consists of a single training session and a subsequent recall test to assess memory formation. While the task is simple, the underlying mechanisms for memory acquisition, consolidation, storage and retrieval are complex. Here, we set up a group of mice which foot shock was delivered immediately after exposed to the IA training chamber to distinguish IA training-induced proteome and phosphoproteomics dynamics from a shock-only stress response. A general question is whether there are proteins which play overlapping or distinctive following IA training and/or immediate shock. GO analysis of proteins regulated by IA training and immediate shock revealed a series of cellular functions that were significantly enriched both in IA and Shock group, or enriched uniquely in either IA or Shock group. Cellular functions, such as positive regulation of adenylate cyclase-activating adrenergic receptor signaling, regulation of post-synapse organization and neuronal synaptic plasticity, regulation of ARF protein signal transduction, protein polymerization, Ca^2+^ transport and actin filament-based process, were found to be enriched in both IA and Shock groups (Figure 4B). We also identified a series of cellular functions that were uniquely enriched in either the IA or Shock group. For example, cellular functions, such as regulation of postsynaptic neurotransmitter receptor activity, SA node cell action potential, release of sequestered Ca^2+^ into cytosol by sarcoplasmic reticulum, synaptic transmission, and anterograde trans-synaptic signaling, were distinctively enriched in IA group. In the same way, we found that cellular functions including positive regulation of LTD, negative regulation of microtubule depolymerization, vesicle docking involved in exocytosis, and regulation of GTPase activity were uniquely enriched in Shock group (Figure 4B). Proteins that exhibit same cellular functions in both IA and Shock groups may be categorized to common proteins that respond to various experiences. On the other hand, proteins linked to cellular functions distinctively enriched in either the IA or Shock group may be categorized by unique proteins that respond differentially to either IA training or immediate shock. These findings support the idea that those behavioral phenotypes elicited by different forms of experience (here IA training and immediate shock) are mediated by proteins or PTMs involved in 1) shared cellular functions that can be regulated in either same or different directions and 2) unique cellular functions that are differentially enriched following specific experiences. Functional validation of PSD proteins or phosphosites that are uniquely regulated by IA training or immediate shock will be required to better understand molecular mechanisms for learning and memory formation.

A central question for all forms of synaptic plasticity is the degree to which phenotypic changes are driven by changes in protein expression and/or PTMs, such as phosphorylation. A previous study indicates a requirement for protein synthesis and enhanced levels of protein phosphorylation for synaptic plasticity (Costa-Mattioli et al., 2009; Montarolo et al., 1986; Woolfrey and Dell’Acqua, 2015). However, the exact time frames that distinguish protein synthesis- or phosphorylation-dependency for learning and memory formation remains unclear (Tully et al., 1994; Villareal et al., 2007). In this study, we analyzed proteins from hippocampal PSD harvested 1hr after IA training or ∼5mins after immediate shock. Interestingly, we observed a decreasing pattern of protein level after IA training (Figure 3B). The degree of decrease was more obvious for phosphorylation level (>50% dephosphorylated) than total protein level (approximately 25%). Methodologically, inhibitory avoidance triggers sequential biochemical reactions in the hippocampus that are important for memory formation, and these biochemical events are similar to those necessary for synaptic plasticity including LTP (Izquierdo and Medina, 1997). LTP is the most studied form of synaptic plasticity and it is the most closely linked molecular mechanism underlying learning and memory. LTP triggers various changes in the postsynaptic sites of neurons including gene expression, neuronal morphology, protein transportation and ion channel properties. Long-term potentiation in the hippocampus is a well-established model for learning and memory (Bliss and Collingridge, 1993; Nicoll, 2017). It was shown that LTP induced by learning *in vivo* mimicked the effects of hippocampal LTP induced by high-frequency stimulation (Asok et al., 2019; Izquierdo et al., 2006; Whitlock et al., 2006). One of the key regulators of these neuronal processes occurring during LTP is protein phosphorylation (Giese and Mizuno, 2013; Lee, 2006). However, our results show overall trend of dephosphorylation in both IA and Shock groups. Because the degree of dephosphorylation is much bigger than reductions in protein level, increased protein phosphatase activity following IA training and immediate shock can be one of the possible mechanisms to explain our results. Among the protein components at the synapse, enzymes controlling protein phosphorylation have been considered important for the induction and maintenance of long-term changes in synaptic strength and, as a counterpart, protein phosphatases have emerged as another key regulator of synaptic plasticity (Coba, 2019; Woolfrey and Dell’Acqua, 2015). We found that Ppm isoforms were regulated differentially by IA training and immediate shock. We were particularly interested in Ppm1h because it has recently been reported that Ppm1h can counteract LRRK2 signaling via Rab protein dephosphorylation, which may potentially link to the molecular mechanisms of LRRK2-mediated neurological disorders such as Parkinson’s disease. We showed that Ppm1h can manipulate levels of glutamate receptors and is affected by neuronal activity both *in vivo* and *in vitro*. The result on Ppm1h functionality is a good example supporting how global dephosphorylation of hippocampal PSD proteome affects IA-mediated learning and memory.

We demonstrated that the PSD proteome underwent dynamic phosphorylation regulation following IA training and immediate shock and this led us to investigate kinases in the PSD (Figure 6). More than 250 kinases are expressed in adult mammalian brains but only a few subsets of kinase, such as calcium/calmodulin-dependent kinase II (CaMKII), extracellular signal regulated kinase 1 and 2 (ERK1/2), cAMP-dependent protein kinase A (PKA), cGMP-dependent protein kinase G (PKG), the phosphatidylinositol 3-kinase (PI3K) and glycogen synthase kinase 3αand 3β(Gsk3α/3β), are known to play critical role in learning and memory (Giese and Mizuno, 2013). We demonstrated that 15 kinases and their phosphorylation sites were dynamically regulated by IA training or immediate shock. It is well known that CaMKIIαis highly expressed in the hippocampus and activated during LTP induction as well as affective learning (Lisman et al., 2002; Lisman et al., 2012). However, previous study showed that CaMKII activation is required during, but not after, training for memory formation by IA training (Murakoshi et al., 2017). Because PSD fractions were prepared 1 hour after IA training, this may explain why IA-trained mice in our study showed no obvious changes compared to the increased level of CaMKIIαin the PSD of mice that received immediate shock. We also found that CaMKIIδ was decreased in IA and further decreased in the shock group. The roles of CaMKIIδ in memory process remain unclear, but there is increasing evidence suggesting that this enzyme can be regulated by training and may contribute to different stages of memory formation. For example, it was found that sustained expression of CaMKIIδ was observed up to 1 week after novel object recognition training and antisense oligo to a CaMKIIδ reversed the effect on memory persistence (Zalcman et al., 2018). In this training paradigm, transcriptional activation via NF-κB and increased histone acetylation in the promoter region of *camk2d* gene resulted in increase of CaMKIIδ expression beyond memory consolidation (Federman et al., 2013). Our results also support the hypothesis that the level of CaMKIIδ could be differentially regulated in different subcellular fractions following different types of behavioral task. We have also demonstrated that a series of kinases and their phosphosites were differentially regulated by IA training or immediate shock (Figure 6). Functions and detailed molecular mechanisms of these kinases will need to be tested.

In this study, we applied iTRAQ and TiSH phosphoproteomics approach to mouse hippocampal PSD fractions and provided a comprehensive proteomic dataset containing hundreds of proteins that showed changes in expression and/or phosphorylation following IA training or immediate shock. We observed a significant decrease of PSD proteome and phosphorylation and the dynamic regulation of synaptic kinases and phosphatases. These results should be interpreted with some caution as we only analyzed PSD samples at the 1 hr post-training time point. Therefore, we do not know how these findings generalize to other post-training timepoints, or whether these phenomena specifically represent the proteome remodeling during early phase of memory formation. Therefore, in future studies it will be interesting to monitor proteome dynamics at multiple time scales to identify key modulators regulating memory formation and its maintenance. In summary, we believe that the dataset from our current study can be used broadly to study the underlying mechanisms for learning and memory formation.

## Acknowledgement

We thank all members of the Huganir laboratory for helpful comments, discussion, and critical reading of the manuscript and especially Drs. Bian Liu, Dylan Hale, Qianwen Zhu and Austin Graves for data curation and critical discussion. We would like to thank especially Ashley Irving, Sarah Rodriguez, Lisa Hamm and Richard Johnson for their technical assistance and administrative support. This work was supported by NIH grants R01 NS036715 and R01 MH112152 (to R.L.H.). This study was supported by the Novo Nordisk Foundation (NNF16OC0023448 to M.R.L.), the Villum Center for Bioanalytical Sciences at SDU (M.R.L.), and the Danish Diabetes Academy (NNF17SA0031406 to T.K.).

## Author Contributions

S.H., T.K., M.R.L. and R.L.H. designed research. S.H. and A.M.B. performed behavioral and biochemical experiments and analyzed data. T.K. performed proteomic analysis and data processing. S.H. and T.K. performed bioinformatic analysis. M.R.L. and R.L.H. supervised the project including funding acquisition and project administration. S.H. T.K., A.M.B. M.R.L. and R.L.H wrote the manuscript.

## STAR★METHODS

Detailed methods are provided in the online version of this paper and include the following:

### • KEY RESOURCES TABLE

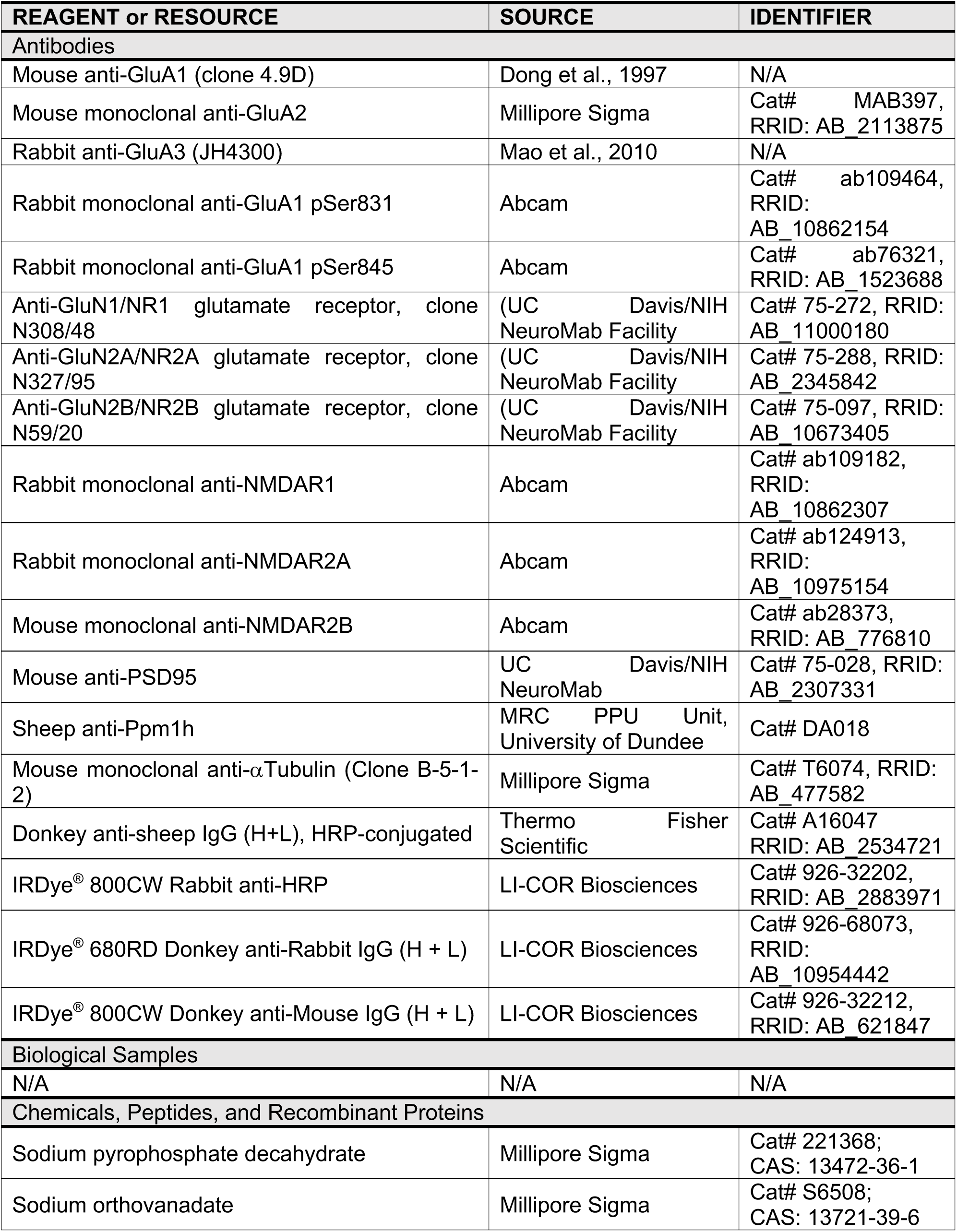
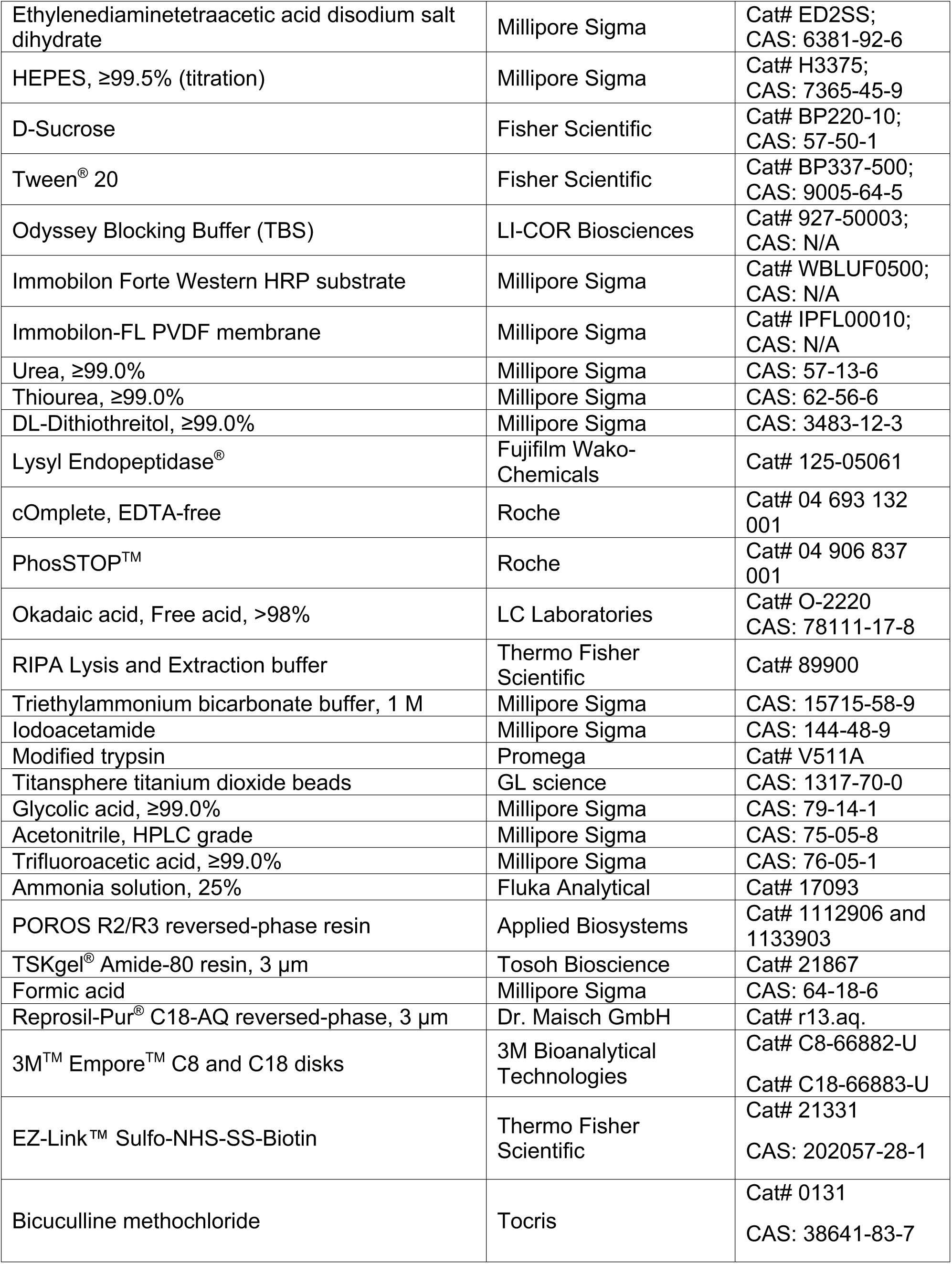
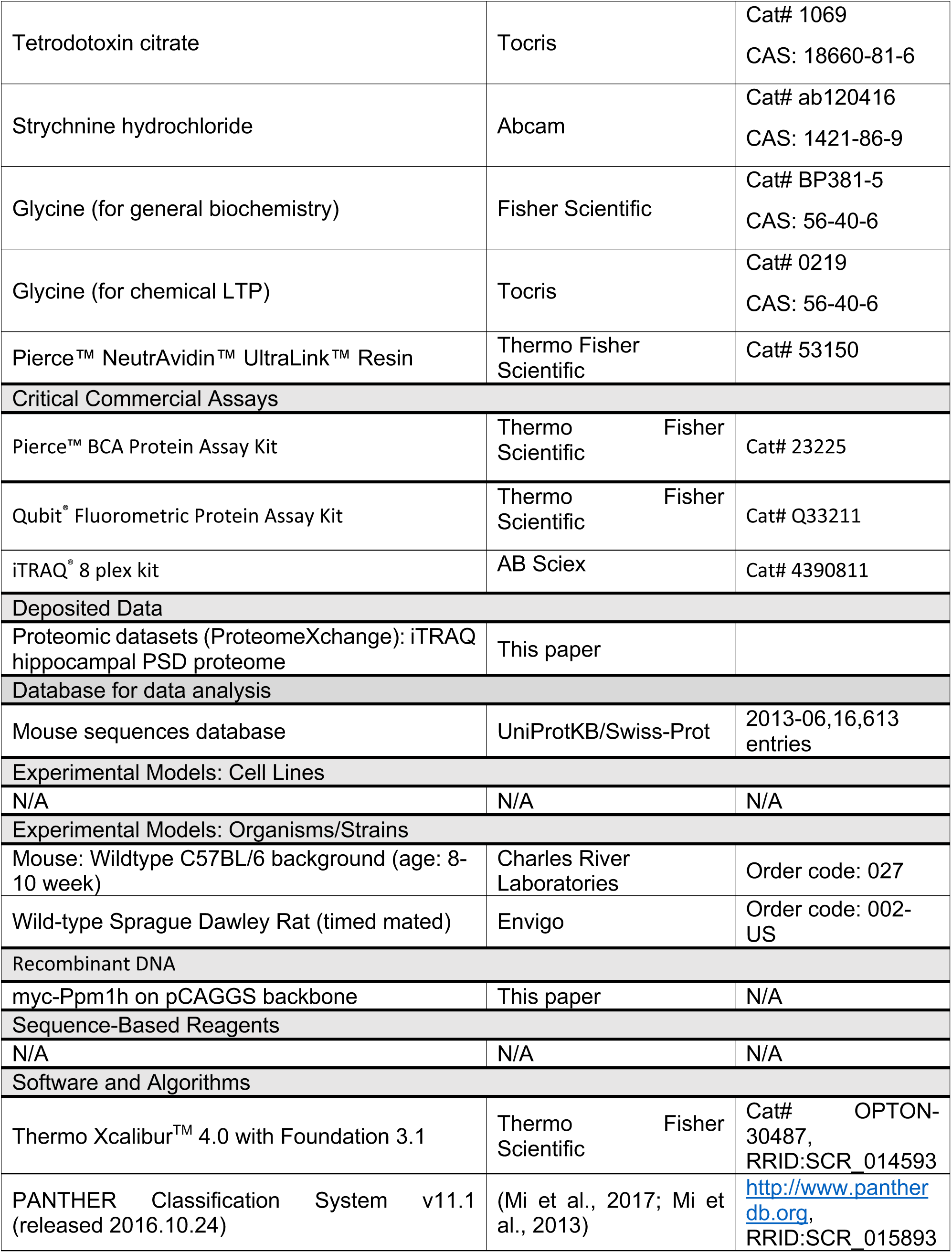
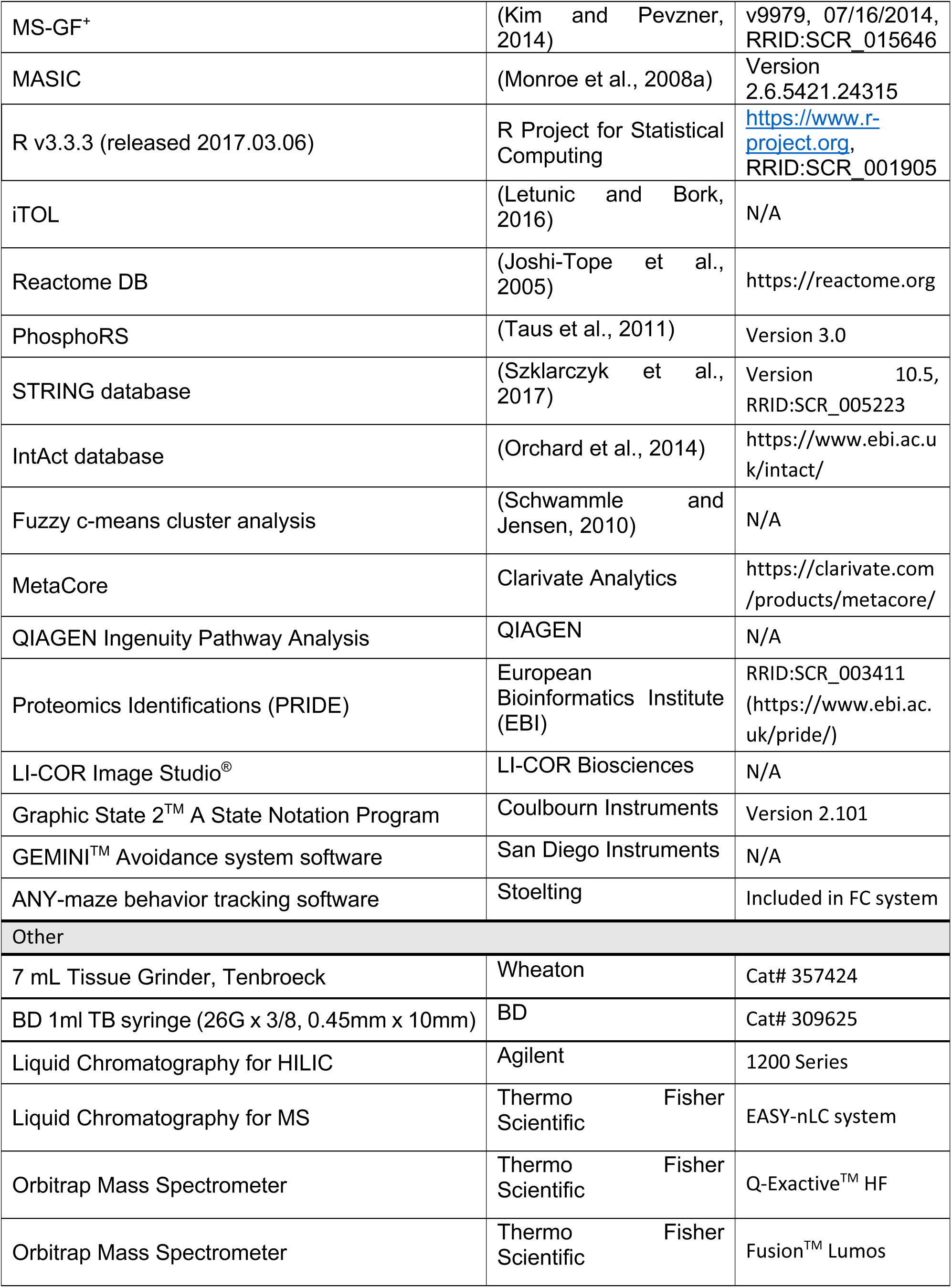
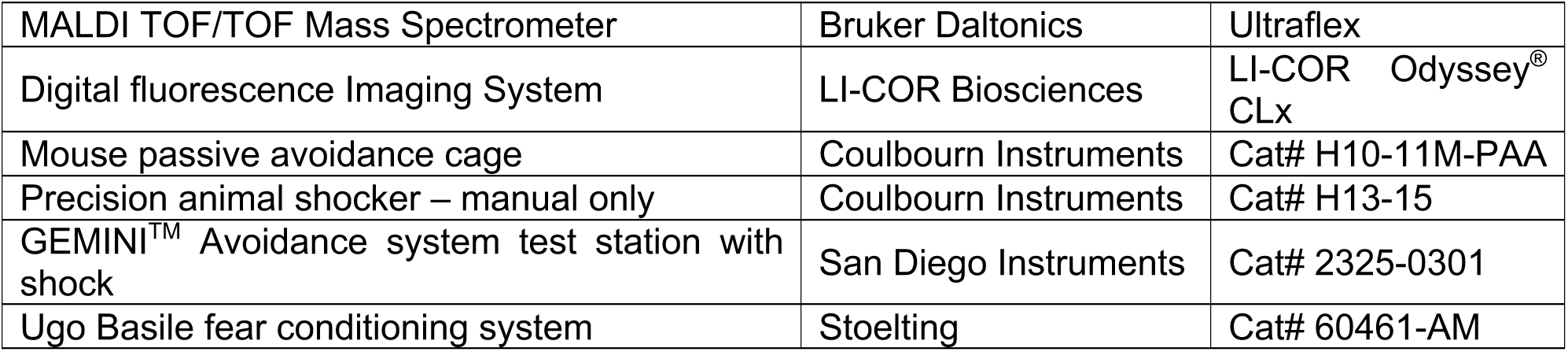

### • CONTACT FOR REAGENT AND RESOURCE SHARING

Further information and requests for resources and reagents should be directed to and will be fulfilled by the Lead Contact, Martin R. Larsen

### • EXPERIMENTAL MODEL AND SUBJECT DETAILS

#### Animal use

All animals were treated in accordance with the Johns Hopkins University Animal Care and Use Committee guidelines. For inhibitory avoidance experiments, mice (purchased from Charles River Laboratories) were delivered at age 8 weeks and group housed for 2 weeks until IA testing. Sprague Dawley rats (purchased from Envigo, former Harlan Laboratories) were used for primary neuronal cultures at embryonic day 18 (E18) as described below. All animals were group housed in a standard 12 hr light/ 12 hr dark cycle. IA testing was conducted during the dark phase.

### • METHOD DETAILS

#### Inhibitory avoidance (IA)

Mice were handled for 5 min on each of the 5 consecutive days before beginning experiments. The inhibitory avoidance (IA) testing cage consisted of a rectangular chamber (35.56 cm wide × 17.78 cm deep × 30.48 cm high, Passive avoidance cage for mouse from Coulbourn Instruments; 24.13 cm wide × 20.32 cm deep × 20.32 cm high, GEMINI^TM^ Avoidance system from San Diego Instruments) divided into two separate compartments, “light” and “dark” compartments. The light compartment was built with transparent Plexiglas and illuminated with a bright overhead stimulus light, while the dark compartment was built with nontransparent Plexiglas and was not illuminated. The compartments were separated by a guillotine door, and both compartments were equipped with metal grid floors connected to an electric generator source that delivered an electric shock (1 mA, 2 seconds). IA testing cage was controlled by Graphic State 2, a state notation program (Coulbourn Instruments). For our experiment, we adopted 3-step IA protocol which consisted of 3 individual sessions, habituation, acquisition, and retention. The latency to enter the dark compartment was recorded as an index of memory consolidation.

For habituation (day 1), a mouse was placed in the light side of the chamber facing the wall of the opposite side of the guillotine door. After 30 seconds the door was opened and the mouse was allowed to explore until it entered the dark compartment. The door closed immediately after the mouse entered the dark side and the mouse was returned promptly to the home cage after entering the dark compartment of the testing cage.

For acquisition (day 2), the mouse again was placed in the light compartment of the testing cage facing the wall of the opposite side of the guillotine door. The door was opened after 30 seconds, and the latency to cross to the dark side following door opening was recorded. The guillotine door closed immediately after the mouse entered the dark compartment, and 3 seconds later the mouse received a foot shock (2 seconds, 1 mA). The mouse remained in the dark chamber for 30 seconds following foot shock for recovery, then it was returned gently to the home cage. Animals in the ‘shock-only’ group were placed in the IA chamber and given a same strength of foot shock (2 seconds, 1 mA) and were immediately removed from the IA chamber for tissue harvest.

For the retention/memory test, 1 hr after training the mouse was reintroduced to the light compartment of the testing cage facing the wall of the opposite side of the guillotine door. The door opened 30 sec after the mouse was placed in the light compartment, and the latency to step through to the dark compartment was recorded as a measure of memory retention (compared with step through latency of acquisition trial). The maximum latency was set at 5 min, after which mice were returned to the homecage. The hippocampus was dissected after completion of the retention test (within 5-10 minutes). Mice were anesthetized with isoflurane for 15 seconds followed immediately by cervical dislocation. Brains were removed and hippocampi were dissected in ice-cold dissection media and immediately frozen with liquid nitrogen. Samples were kept at -80°C until subcellular fractionation for PSD preparation.

#### Enriched environment (EE)

For all experiments mice were age 8-10 weeks. Mice were first handled and habituated to minimize stress-induced changes. Mice were then either left in their home cage (control) or allowed to explore an enriched environment (EE) which is composed of novel objects, tubes, and strings of beads suspended from the cage lid in a large cage for 8 hours during wake period then transferred back to the home cage. This is a physiologically relevant condition which expect to drive neuronal activity and synaptic plasticity (Nithianantharajah and Hannan, 2006; Rampon et al., 2000). On the next day, mice were exposed to the EE cage with different sets of novel objects in a different arrangement for 8 hours. This 2-day EE cycle was repeated for 2 weeks. On the last day, mice were exposed to the EE chamber then anesthetized by inhalation of isoflurane for 15 seconds followed immediately by cervical dislocation. Brains were removed and hippocampi were dissected in ice-cold dissection media and immediately frozen with liquid nitrogen. Samples were kept at -80°C until subcellular fractionation for PSD preparation.

#### Dissociated rat neuronal culture

Cortical neurons obtained from pregnant wild-type Sprague Dawley rats (purchased from Envigo) at embryonic day 18 were initially prepared in Neurobasal media (Invitrogen) supplemented with 2% B-27, 2 mM GlutaMax, 50 U/mL penicillin, 50 mg/mL streptomycin, and 5% horse serum (Invitrogen) and plated onto poly-L-lysine-coated tissue culture dishes at a density of 800,000 cells per well. Cortical neurons were then transferred and maintained in a humidified tissue culture incubator at 37 °C in a 95% air and 5% CO_2_ mixture; 5 mM FDU (5-Fluoro-2′-Deoxyuridine and 5 mM Uridine; Sigma) was added at DIV4 to inhibit glia proliferation and cells were thereafter maintained in NM1 (Neurobasal media with 2% B-27, 2 mM GlutaMax, 50 U/mL penicillin, 50 mg/mL streptomycin, and 1% horse serum). Cultured cortical neurons were fed twice per week. Cortical neurons were grown for 11–12 days in vitro for induction of homeostatic up- and down-scaling and 18-19 days in vitro for induction of chemical LTP. For ppm1h overexpression experiments, cortical neurons were electroporated with myc-ppm1h construct at DIV0 using Rat Neuron Nucleofector kit (Lonza) following manufacturer’s manual, and cells were used when 2-3 weeks old.

For induction of homeostatic up- or down-scaling, cortical neurons (DIV11-12) were treated with bicuculline (20 µM) or TTX for 48 hrs followed by either direct lysis in RIPA buffer containing protease inhibitor cocktail (Roche), phosphatase inhibitor cocktail (Roche) and 1 µM okadaic acid, or surface biotinylation. For glycine-induced chemical LTP experiments, cortical neurons (DIV19-20) were first preincubated with Mg^2+^-ACSF (143 mM NaCl, 5 mM KCl, 10 mM HEPES [pH 7.42], 10 mM Glucose, 2 mM CaCl_2_, 1 mM MgCl_2_, 0.5 µM TTX, 1 µM Strychnine, and 20 µM BIC), followed by glycine treatment for 10 min (chemical LTP ACSF: 200 µM glycine / 0 Mg^2+^), and returned to the original Mg^2+^-ACSF (0 glycine / 1 mM MgCl_2_) for 30 min prior to surface biotinylation/lysis.

#### Surface biotinylation

Neurons were rinsed with ice-cold PBS containing 0.1 mM CaCl_2_ and 1 mM MgCl_2_ (pH 8.0) (PBS-CM), then incubated in PBS-CM containing 1 mg/ml Sulfo-NHS-SS-biotin (Thermo Fisher Scientific, 30 min, 4°C). After biotinylation reaction, neurons were rinsed with PBS-CM, and the biotinylation reaction was quenched in PBS-CM containing 50 mM glycine (2 x 5 min, 4°C). Cells were lysed in RIPA buffer containing protease inhibitor cocktail (Roche), phosphatase inhibitor cocktail (Roche) and 1 µM okadaic acid, then cleared by centrifugation (17,000 x g, 10 min, 4°C). Protein concentration of each lysate was quantified using BCA protein assay kit (Thermo Fisher Scientific), and equal amounts of protein were incubated overnight with NeutrAvidin-coupled agarose beads (Thermo Fisher Scientific) at 4°C with gentle rotation. Beads were washed three times with ice-cold lysis buffer, and biotinylated proteins were eluted with 2x SDS sample buffer. Cell-surface or total proteins were then subjected to SDS-PAGE and analyzed by Western blot.

#### Sub-cellular fractionation and Western blotting

For post-synaptic density preparation, hippocampi dissected immediately following memory recall (one-hour following IA-training) were homogenized using 20 strokes from syringes equipped with 26G x 3/8 (0.45mm x 10mm) needles in homogenization buffer (320 mM sucrose, 5 mM sodium pyrophosphate, 1 mM EDTA, 10mM HEPES pH 7.4, 200 nM okadaic acid, 1 mM sodium orthovanadate, protease inhibitor cocktail (Roche), phosphatase inhibitor cocktail (Sigma-Aldrich)). The homogenate was then centrifuged at 1,000 x g for 10 minutes at 4°C to yield P1 (nuclear fraction) and post-nuclear supernatant (PNS) fractions. PNS fraction was further centrifuged at 17,000 x g for 20 minutes at 4°C to yield P2 (membrane/crude synaptosome) and S2 (cytosol) fractions. P2 was resuspended in hypotonic resuspension buffer (Milli-Q^®^ water with 5 mM sodium pyrophosphate, 1 mM EDTA, 10mM HEPES pH 7.4, 200 nM okadaic acid, 1 mM sodium orthovanadate, protease inhibitor cocktail (Roche), phosphatase inhibitor cocktail Roche)), then centrifuged at 25,000 x g for 20 minutes at 4°C to yield lysed synaptosome (LS) fractions. Collected LS fractions were resuspended in resuspension buffer (50 mM HEPES pH 7.4, 5 mM sodium pyrophosphate, 1 mM EDTA, 200 nM okadaic acid, 1 mM sodium orthovanadate, protease inhibitor cocktail (Roche), phosphatase inhibitor cocktail (Roche)) and then mixed with an equal part of 1% Triton X-100 (containing protease and phosphatase inhibitors). This mixture was incubated at 4°C with rotation for 10 minutes followed by centrifugation at 50, 000x g for 20 minutes at 4°C to yield PSD preparation. The final PSD pellet was resuspended in 50 mM HEPES pH 7.4 (containing protease and phosphatase inhibitors). The protein concentration from PSD fractions was determined using BCA protein assay followed by biochemical analysis.

For Western blotting analysis, samples were quantified using BCA protein assay kit and loaded onto 9 or 12% SDS-PAGE (depending on the molecular weights of the protein of interest). Proteins were transferred to PVDF membrane, and the membranes were blocked with Odyssey blocking buffer for fluorescent detection for 1 hr at room temperature. Primary antibodies were resuspended in Odyssey blocker / TBS-T (1X TBS supplemented with 0.2% Tween^®^ 20) mixture (Odyssey blocker : TBS-T = 1 : 1) and incubated overnight at 4°C with gentle rocking. Primary antibodies were removed and membranes were washed followed by IRDye^®^-conjugated secondary antibody incubation in blocking solutions. For primary antibodies where IRDye^®^- conjugated secondary antibodies were not available, membranes were first probed with HRP (horseradish peroxidase)-conjugated secondary antibodies followed by re-probing with IRDye^®^- conjugated anti-HRP antibody. Blots were developed using either LI-COR Odyssey^®^ CLx Imaging system (LI-COR).

#### Sample preparation for mass spectrometry analysis

##### In-solution trypsin and Lys-C digestion

PSD fractions isolated from mouse hippocampi were lysed, reduced, and predigested in 6 M Urea, 2 M Thiourea, containing 10 mM Dithiothreitol and 2 µl Lys-C endopeptidase supplemented with PhosSTOP^TM^ phosphatase inhibitor for 2 h at room temperature (RT). Thereafter, the lysates were diluted 10 times using 20 mM Triethylammonium bicarbonate buffer (TEAB; pH adjusted to 7.5) and tip-sonicated for 2 x 20 seconds on ice. Samples were then alkylated by 20 mM iodoacetamide for 20 min in the dark before digestion with 2% (w/w) trypsin overnight at 37°C.

##### iTRAQ labeling of peptides

Peptide concentration was measured by Qubit^®^ Fluorometric protein assay according to the manufacturer’s instructions. A total of 60 µg was aliquoted from all samples (4 groups from total lysates and PSD fractions) and lyophilized before labeling with iTRAQ^®^ 8 plex kit (AB Sciex). Three biological replicates were made and labeling was performed as follows: total naïve 113, total walk-through 114, total shock-only 115, total IA-trained 116, PSD naïve 117, PSD walk-through 118, PSD shock-only 119, PSD IA-trained 121. The labeling was performed according to the manufacturer’s protocol and complete labeling was validated by running combined aliquots on MALDI MS (Bruker Daltonics, Germany). The equal amount (60 µg) of protein per sample were mixed in equal ratios and stored at -20°C until phosphopeptide enrichment.

##### Enrichment of phosphorylated peptides

The purification of phosphopeptides was performed according to a slightly modified TiSH (TiO2-SIMAC-HILIC) phosphopeptide enrichment procedures (Engholm-Keller and Larsen, 2016; Kang et al., 2018; Thingholm et al., 2006), in which nonmodified peptides are first separated from phosphopeptide species using TiO_2_ beads. Briefly, the lyophilized iTRAQ labeled sample was made up to 1 ml loading buffer [1 M glycolic acid, 80% acetonitrile (ACN), 5% TFA] and added with TiO_2_ beads at 0.6 mg/100 µg (bead/peptide), and incubated at RT for 10 min. The suspension was centrifuged for 15 sec in a table centrifuge and the supernatant loaded onto a second batch of TiO_2_ (containing half the amount of TiO_2_ as initially used) and incubated at RT for 15 min. The two batches of TiO_2_ were washed with 100 µl of washing buffer 1 [80% ACN, 1% trifluoroacetic acid (TFA)] and centrifuged for 15 sec in a tabletop centrifuge. The supernatant was removed, and the beads were washed with 100 µl washing buffer 2 (10% ACN, 0.1% TFA) and centrifuged for 15 sec in a tabletop centrifuge. The supernatant was removed, and the beads were dried in a vacuum centrifuge for 5 min. The bound peptides were eluted with 100 µl of 1% ammonium hydroxide for 15 min and then centrifuged at 1,000 g for 1 min. The eluted peptides were passed over a C8 stage tip (Thingholm and Larsen, 2009) to retain the TiO_2_ beads and dried by vacuum centrifugation to produce the enriched phosphopeptide fraction. The flow through from the initial loading buffer (containing nonmodified peptides) and washes were combined and dried by vacuum centrifugation to produce the nonmodified peptide fraction. The nonmodified peptide fraction was acidified with TFA and desalted on a R3 stage tip column before HILIC fractionation.

##### Sample desalting

Samples were desalted before HILIC fractionation. The desalting columns were self-made by inserting a small plug of C18 material into the constricted end of a 200 µl tip and packed with a mixture of R2 and R3 reversed-phase resin applying manual air pressure with a syringe, followed by an optimized desalting procedure (Thingholm et al., 2006). Briefly, the samples were acidified before loading onto the columns (equilibrated with 0.1% TFA), followed by washing with 0.1% TFA, and peptides were eluted using 60% ACN, 0.1% TFA and were lyophilized before further processing.

##### Hydrophilic Interaction Liquid Chromatography (HILIC)

The phosphorylated and the nonmodified peptide samples were subjected to fractionation using HILIC (Kang et al., 2018). Briefly, these samples were resuspended in 90% ACN, 0.1% TFA (Solvent B) and loaded onto a 450 µM OD x 320 µM ID x 17 cm micro-capillary column packed with TSKgel^®^ Amide-80 resin material using an Agilent 1200 Series HPLC. Peptides were separated using a gradient from 100–60% Solvent B (Solvent A: 0.1% TFA) running for 30 min at a flow-rate of 6 µl/min. The fractions were automatically collected in a 96 well plate at one-minute intervals after UV detection at 210 nm.

#### LC-MS/MS, proteomic data handling and bioinformatic analysis

##### Reverse-phase nanoLC-ESI-MS/MS analysis

All fractions were redissolved in buffer A (0.1% FA) and analyzed using a nLC-MS/MS system consisting of an Easy-nLC and an Orbitrap Fusion^TM^ Lumos (phospho-proteome) or a Q-exactive^TM^ HF (proteome) mass spectrometers (MS) were used separately to increase the speed of analysis. The samples were loaded onto a 2 cm pre-column (100 μm inner diameter) and separated on a 17 cm fused silica capillary column (75 μm inner diameter). All columns were homemade and packed with ReproSil-Pur^®^ C18 AQ 3 μm reversed-phase resin material. The peptides were eluted using 73–133 min gradients from 1 to 40% buffer B (95% ACN, 0.1% FA) and introduced into the MS instrument via nanoelectrospray according to the intensity of each HILIC peptide fraction. A full MS scan in the mass area of 400–1400 Da was performed in the Orbitrap with a resolution of 120,000, an AGC target value of 5 × 10^5^, and a maximum injection time of 100 ms. For each full scan, ‘Top speed’ mode was selected for higher energy collision dissociation (HCD). The settings for the HCD were as follows: AGC target value of 3 × 10^4^, maximum injection time of 60 ms, isolation window of 1.2 Da, and normalized collision energy of 38. All raw data were viewed in Xcalibur^TM^ v4.0.

##### MS Data processing and statistical analysis

The raw MS data sets were processed for protein/peptide identification using the MS-GF+ (v9979, 07/16/2014) (Kim and Pevzner, 2014) combined MASIC (Monroe et al., 2008b) pipeline with a peptide mass tolerance of 20 ppm, reporter ion m/z tolerance half width of 2 mDa, and a false discovery rate (FDR) of 1% for proteins and peptides. All peak lists were searched against the UniProtKB/Swiss-Prot database of mouse sequences (06/2013, 16,613 entries) with decoy using the parameters as follows: enzyme, trypsin; maximum missed cleavages, 2; fixed modification, carbamidomethylation (C), iTRAQ tags (K, peptide N termini); variable modifications, oxidation (M) and phosphorylation (S, T, Y). Data sets with raw MS values were filtered to remove potential errors using several criteria. For relative protein quantification, the output tsv file from MS-GF^+^ combine MASIC pipeline was imported into Microsoft Excel and filtered as follows: elimination of contaminants and reversed sequences for each accession number. Protein relative expression values from the respective unique or razor peptides were calculated by summing all unique/razor peptides intensity of each protein and normalized to the number of total intensities of each group estimating the relative amounts of the different protein within the relative sample. The resulting ratios were logarithmized (base = 2) to achieve a normal distribution. Ratios were averaged over overlapping proteins or phosphopeptides. Fragment ion masses (b- and y-type ions) with the phosphorylation were checked at the peptide backbone in the MS/MS data sets using PhosphoRS. Fuzzy c-means cluster analysis was performed by using expression changes for regulated proteins/phosphopeptides.

##### Bioinformatic processing and data analysis

Gene Ontology annotation enrichment analysis was performed using the PANTHER. Reactome pathway analysis was used to functionally annotate genes implicated in canonical pathways, using an FDR threshold of 0.05. The regulated proteins and phosphorylation were searched against the STRING database (version 10.5) and IntAct database (https://www.ebi.ac.uk/intact/) for protein-protein interactions. Ingenuity Pathway Analysis (IPA; Ingenuity Systems) was used to functionally annotate genes implicated in causal biological pathways and functions.

We used the FASTA sequences of the kinase domains retrieved from the KinBase resource or phosphatase domains from the Unitprot database and aligned them by ClustalX2.1 using default parameters for multiple alignment and bootstrapping N-J tree. Kinase or phosphatase sequences were visualized by phylogenetic distances using the Interactive Tree of Life (ITOL) tool (https://itol.embl.de).

### QUANTIFICATION AND STATISTICAL ANALYSIS

Significantly regulated proteins or phosphopeptides were accepted if the z-test for adjusted *p*-value was < 0.05 (95% confidence) with the Benjamini and Hochberg correction in a normal distribution or if the corrected *p*-value was < 0.1 (90% confidence) with a coefficient of variation (CV%) of 30% or smaller. Regulated proteins or phosphopeptides had the same ratio direction (log_2_ ratio, positive or negative) in all of identified biological replicates.

### DATA AND SOFTWARE AVAILABILITY

The proteomics data and search results associated with this study have been deposited to the ProteomeXchange Consortium (Deutsch et al., 2020) via the PRIDE (Perez-Riverol et al., 2019) partner repository.

### ADDITIONAL RESOURCES

N/A

## Acknowledgements

We thank members of the R.L.H. laboratory for helpful comments, discussion and critical reading of the manuscript. This work was supported by grants from the National Institute of Health Grants (R01NS036715, R01MH112152, to R.L.H.), the Novo Nordisk Foundation (NNF16OC0023448 to M.R.L), and the Villum Center for Bioanalytical Sciences at SDU (M.R.L), and the Danish Diabetes Academy (NNF17SA0031406 to T.K.).

---

**Figure S1.**
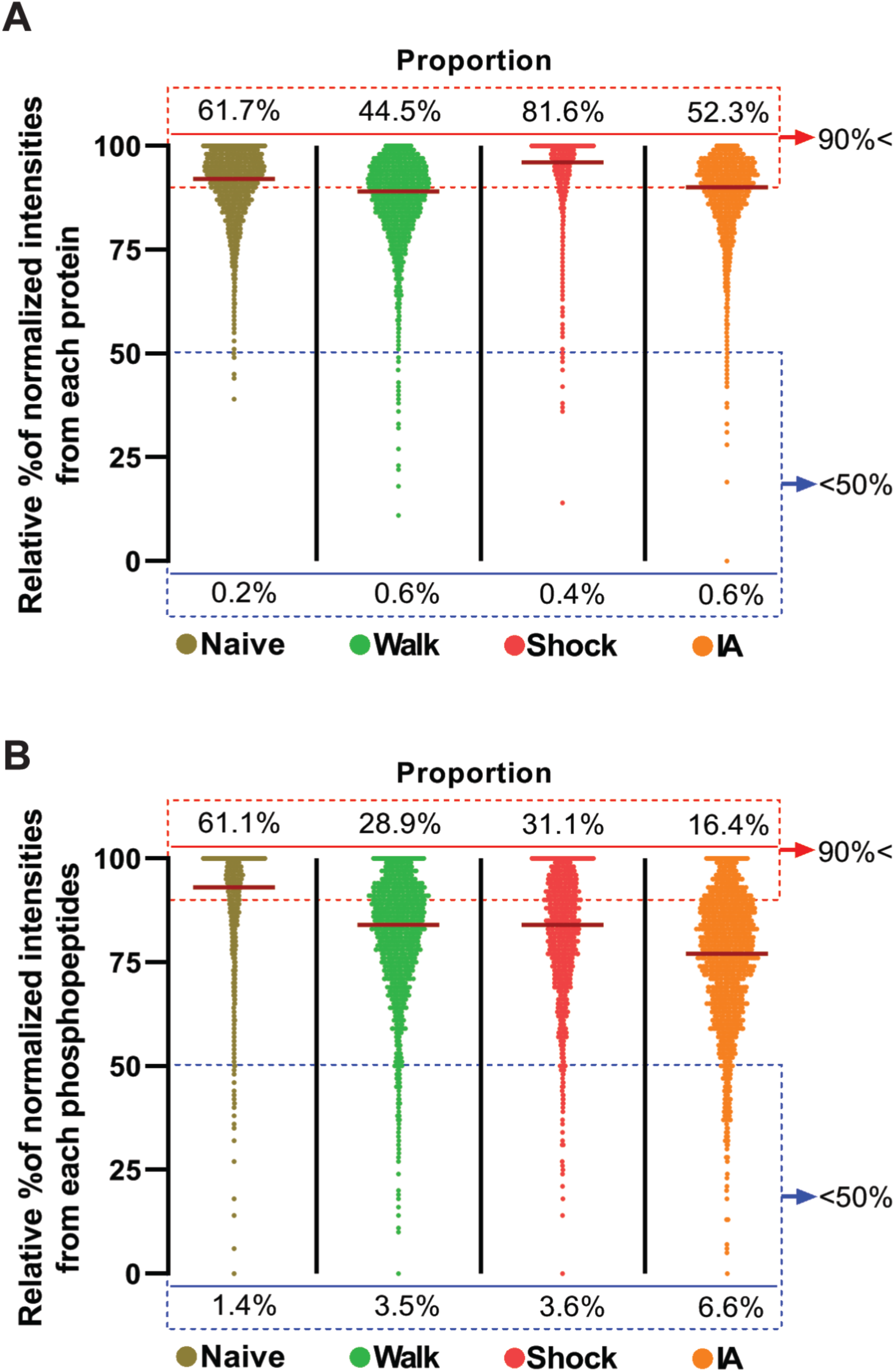
Dynamic changes of the proportion of PSD proteins and phosphoproteins following IA-training and immediate shock. (Related to Figure 3B) Violin plots showing relative percentile of normalized intensities of all identified and quantified (A) proteins and (B) phosphopeptides in the PSD. Y-axis represents proportions of relative (%) intensities of proteins and phosphopeptides per group. Dark red lines inside of the violin plots indicate the median of the relative percentiles. The width of the plots represents the density of proteins and phosphopeptides. Percentile values in the box with red dashed line indicate the proportion values (%) of the proteins and phosphopeptides which have relative percentile of normalized intensities higher than 90%. Percentile values in the box with blue dashed line indicate the proportion values (%) of the proteins and phosphopeptides which have relative percentile of normalized intensities lower than 50%. This result demonstrates that the overall regulation of PSD proteome shows a more obvious decreasing pattern in phosphorylation levels (proportion above 90% intensities: Naïve > Shock > Walk > IA, proportion below 50% intensities: Naïve < Walk < Shock < IA) than non-modified protein levels following IA-training or immediate shock.

**Figure S2.**
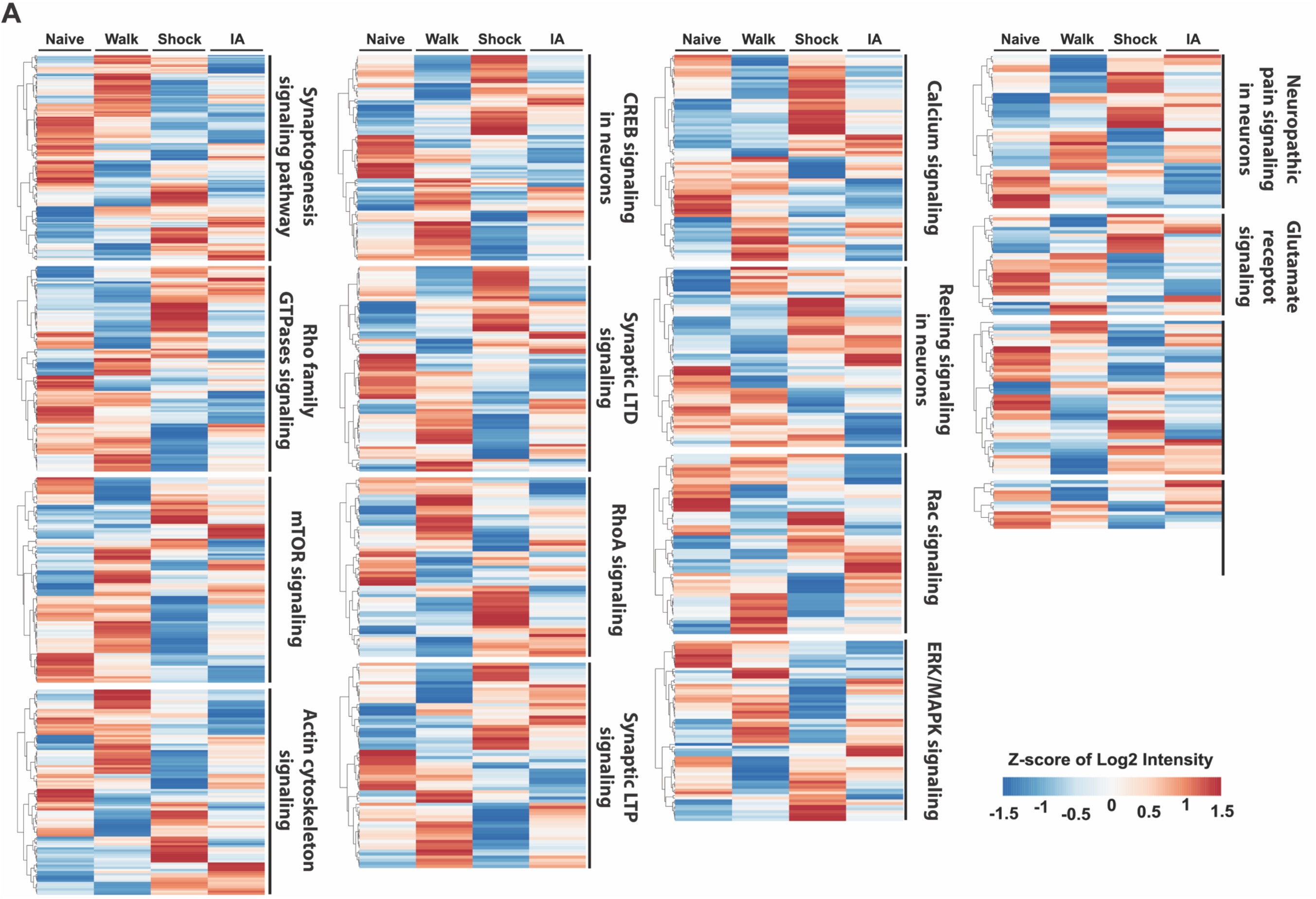
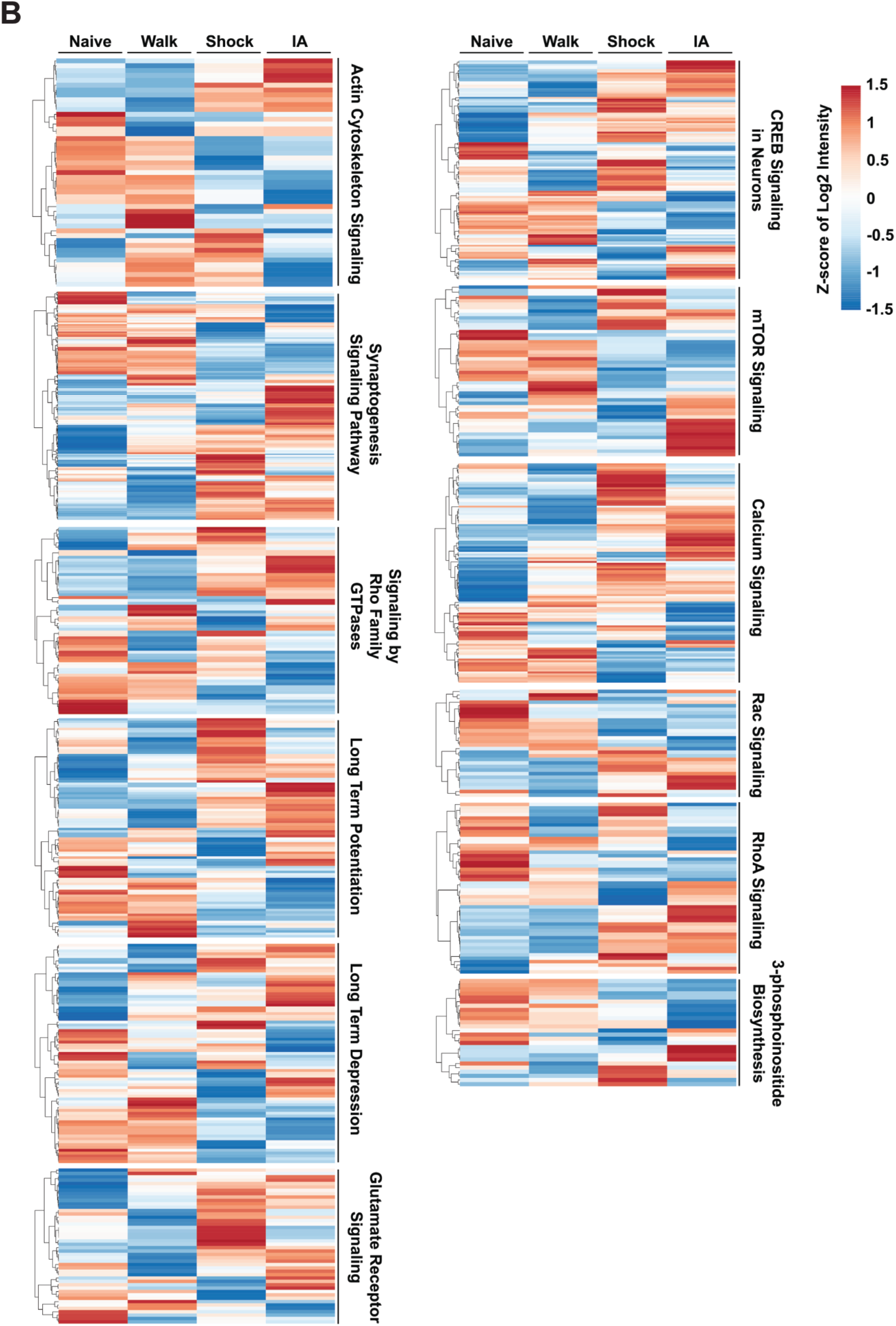
Experience-dependent regulation of PSD proteome and phosphoproteome and related signaling pathways. (Related to Figure 5) (A) Heatmap ordered by hierarchical clustering shows the experience-dependent dynamic regulation of PSD proteins that are involved in the same signaling pathways (1% FDR). (B) Heatmap ordered by hierarchical clustering shows the experience-dependent dynamic regulation of PSD phosphoproteins that are involved in the same signaling pathways (1% FDR). Values for individual protein (row) from all groups (column) are color-coded based on the z-scores of Log_2_ transformed intensity of proteins. These results indicate that proteins and/or phosphoproteins in the PSD that are involved in the same signaling pathways exhibit different roles following different types of experiences.

**Figure S3.**
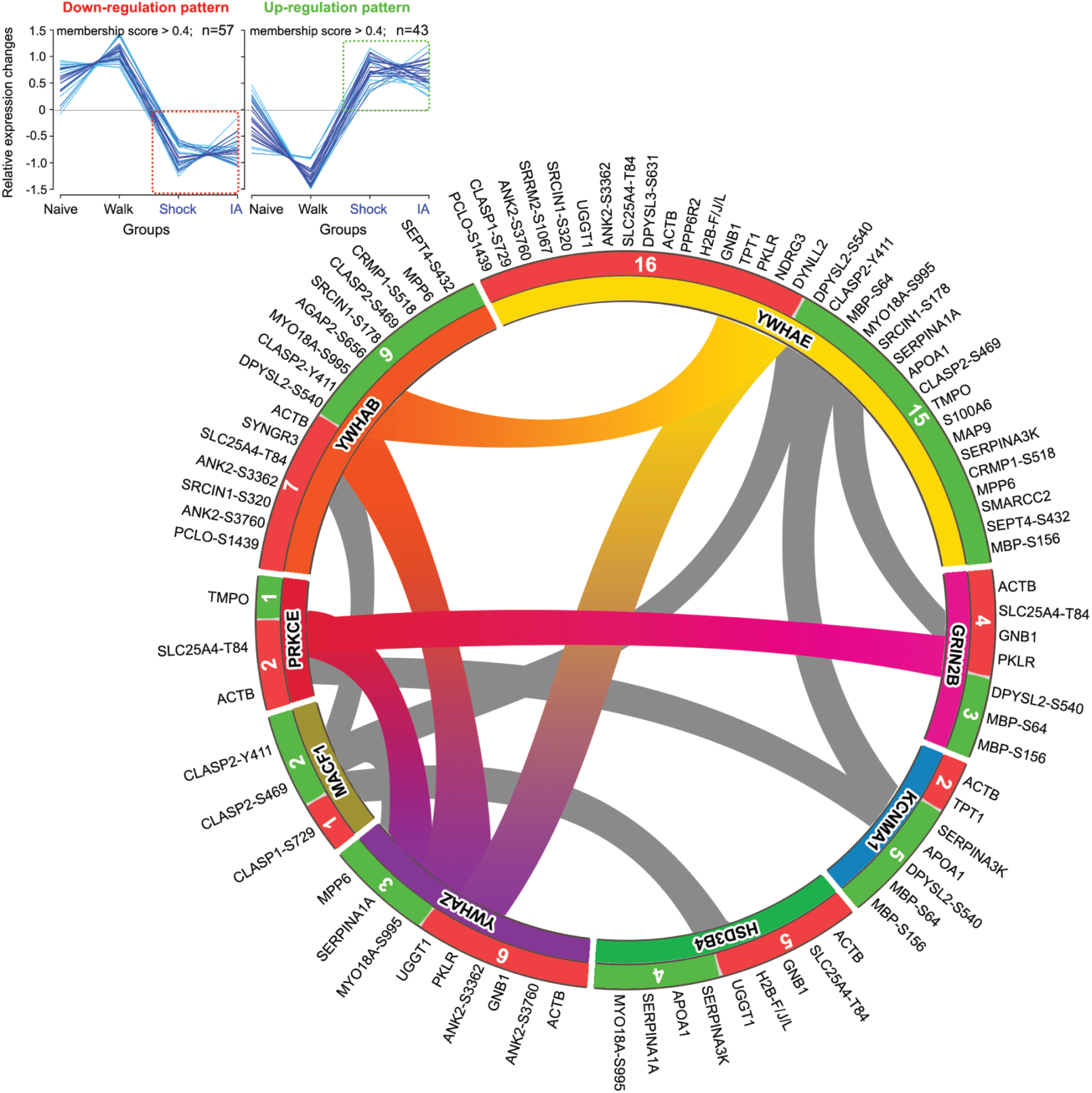
Protein interaction networks of the regulated PSD proteins and phosphoproteins by both IA-training and immediate shock. (Related to Figure 5) Protein interaction map showing upstream proteins and their interacting proteins generated using combinatorial resources (IntAct, STRING, DAVID, Pubmed, and UniProt) and Fuzzy c-means clustering analysis, indicating significantly modulated proteins and phosphoproteins that show up- or down-regulation following both IA-training and immediate shock. Clustering analysis (membership > 0.4) showing up- (n = 43) or down-regulated (n = 57) proteins and phosphoproteins in both IA and Shock groups are shown in the upper left panel. The inner rim of the circular interactome map (CIM) represents possible upstream modulators (*p*-value < 0.05) and the outer rim represents their interacting proteins. The red-colored outer rim (including the number of proteins and phosphoproteins assigned to the classified upstream proteins) of the CIM indicates the down-regulated upstream interacting proteins and phosphoproteins following both IA-training and immediate shock (down-regulation pattern; left). The green-colored outer rim (including the number of proteins and phosphoproteins assigned to the classified upstream proteins) of the CIM indicates the up-regulated upstream interacting proteins and phosphoproteins following both IA-training and immediate shock (up-regulation pattern; right). The *p*-values for individual upstream modulator are listed below: 14-3-3 protein beta, YWHAB (*p*-value: 0.00002); 14-3-3 protein epsilon, YWHAE (*p*-value: 0.001); GRIN2B (*p*-value: 0.002); calcium-activated potassium channel subunit alpha-1, KCNMA1 (*p*-value: 0.002); 3 beta-hydroxysteroid dehydrogenase type 4, HSD3B4 (*p*-value: 0.03); 14-3-3 protein zeta, YWHAZ (*p*-value: 0.04); microtubule-actin cross-linking factor 1, MACF1 (*p*-value: 0.05); protein kinase C epsilon type, PRKCE (*p*-value: 0.09).

**Figure S4.**
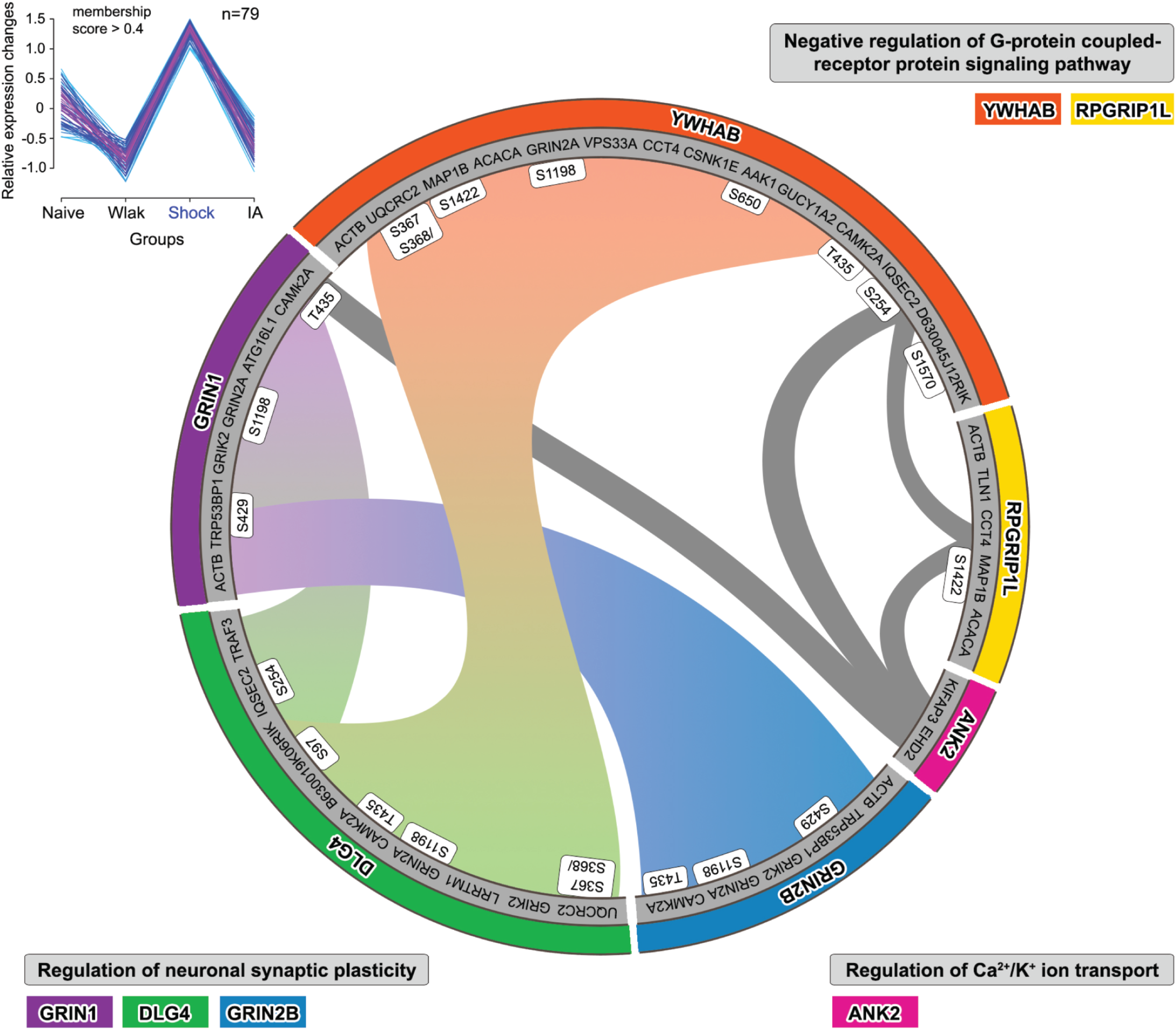
Protein interaction networks of up-regulated proteins and phosphoproteins by immediate shock. (Related to Figure 5) Protein interaction networks of upstream proteins and their interacting proteins associated with distinctively up-regulated proteins and phosphoproteins (with their relevant phosphosites) following immediate shock, generated using combinatorial resources (IntAct, STRING, DAVID, Pubmed, and UniProt) and Fuzzy c-means clustering analysis. Clustering analysis (membership > 0.4) showing up-regulated proteins and phosphoproteins (n = 79) in the Shock group is shown in the upper left panel. The inner rim of the CIM represents interacting proteins and outer rim represent upstream proteins. PSD proteins and phosphoproteins within this cluster reveal 6 regulated upstream proteins that belong to 3 functional annotations (gray box outside of the CIM). The *p*-values for individual parent regulator are listed below: 14-3-3 protein beta, YWHAB (p-value: 0.0001); RPGR-interacting protein 1-like protein, RPGRIP1L (p-value: 0.07); ankyrin 2, ANK2 (p-value: 0.06); glutamate receptor NMDA2B, GRIN2B (p-value: 0.02); disks large homolog 4, Dlg4 (p-value: 0.002); glutamate receptor NMDA1, GRIN1 (p-value: 0.0006).

## SUPPLEMENTAL TABLES

**Table S1 (separate file, related to Figure 2). List of PSD proteins and phosphoproteins (including phosphopeptides and phosphosites) that were identified and quantified from this study (identified and quantified from at least 2 out of 3 biological replicates).**

PSD proteins and phosphoproteins (identified and quantified from at least 2 out of 3 biological replicates from this study) are listed. Sheet #1 shows description on columns of each table. Sheet #2 (Proteome) shows the list of PSD proteins and Sheet #3 (Phosphoproteome) shows the list of phosphoproteins including the information on phosphopeptides and phosphosites.

**Table S2 (separate file, related to Figure 4B). List of proteins and phosphoproteins from mouse hippocampal PSD fractions distinctively regulated by IA-training (IA-unique) or immediate shock (Shock-unique).**

Sheet #1 shows description on columns of each table. Sheet #2 (Proteome) and Sheet #3 (Phosphoproteome) show the list of hippocampal PSD proteins and phosphoproteins (including phosphopeptides and phosphosites information) distinctively regulated by IA training (Figure 4B left panel) or immediate shock (Figure 4B right panel).

**Table S3 (separate file, related to Figure 4C). List of proteins and phosphoproteins from mouse hippocampal PSD fractions regulated same-directionally or bi-directionally by IA-training or immediate shock.**

Sheet #1 shows description on columns of each table. Sheet #2 and #4 (#2: PROT_same-directional, #4: PHOS_same-directional, related to Figure 4C left panel) show the list of hippocampal PSD proteins and phosphoproteins (including phosphopeptides and phosphosites information) that were same-directionally regulated by IA-training or immediate shock. Sheet #3 and #5 (#3: PROT_bi-directional, #4: PHOS_bi-directional, related to Figure 4C right panel) show the list of PSD proteins and phosphoproteins (including phosphopeptides and phosphosites information) that were bi-directionally regulated by IA-training or immediate shock.

**Table S4 (separate file, related to Figure S2). List of regulated proteins and phosphoproteins from mouse hippocampal PSD fractions involved in distinct signaling pathways shown in Figure S2.**

Sheet #1 shows description on columns of each table. Sheet #2 and #3 (#2: PROT_Shock-induced, #3: PROT_IA-induced, related to Figure S2A) show the list of hippocampal PSD proteins that were regulated by immediate shock and IA-training. Sheet #4 and #5 (#4: PHOS_Shock-induced, #5: PHOS_IA-induced, related to Figure S2B) show the list of hippocampal PSD phosphoproteins that were regulated by IA-training and immediate shock.

## Reference List

Asok, A., Leroy, F., Rayman, J.B., and Kandel, E.R. (2019). Molecular Mechanisms of the Memory Trace. Trends Neurosci 42, 14–22.

Bayes, A., and Grant, S.G. (2009). Neuroproteomics: understanding the molecular organization and complexity of the brain. Nat Rev Neurosci 10, 635–646.

Best, P.J., and Orr, J., Jr. (1973). Effects of hippocampal lesions on passive avoidance and taste aversion conditioning. Physiol Behav 10, 193–196.

Bevilaqua, L.R., Medina, J.H., Izquierdo, I., and Cammarota, M. (2005). Memory consolidation induces N-methyl-D-aspartic acid-receptor- and Ca2+/calmodulin-dependent protein kinase II-dependent modifications in alpha-amino-3-hydroxy-5-methylisoxazole-4-propionic acid receptor properties. Neuroscience 136, 397–403.

Bliim, N., Leshchyns’ka, I., Keable, R., Chen, B.J., Curry-Hyde, A., Gray, L., Sytnyk, V., and Janitz, M. (2019). Early transcriptome changes in response to chemical long-term potentiation induced via activation of synaptic NMDA receptors in mouse hippocampal neurons. Genomics 111, 1676–1686.

Bliss, T.V., and Collingridge, G.L. (1993). A synaptic model of memory: long-term potentiation in the hippocampus. Nature 361, 31–39.

Borovok, N., Nesher, E., Levin, Y., Reichenstein, M., Pinhasov, A., and Michaelevski, I. (2016). Dynamics of Hippocampal Protein Expression During Long-term Spatial Memory Formation. Mol Cell Proteomics 15, 523–541.

Brown, R.E., and Milner, P.M. (2003). The legacy of Donald O. Hebb: more than the Hebb synapse. Nat Rev Neurosci 4, 1013–1019.

Cammarota, M., Bernabeu, R., Levi De Stein, M., Izquierdo, I., and Medina, J.H. (1998). Learning-specific, time-dependent increases in hippocampal Ca2+/calmodulin-dependent protein kinase II activity and AMPA GluR1 subunit immunoreactivity. Eur J Neurosci 10, 2669–2676.

Cammarota, M., Izquierdo, I., Wolfman, C., Levi de Stein, M., Bernabeu, R., Jerusalinsky, D., and Medina, J.H. (1995). Inhibitory avoidance training induces rapid and selective changes in 3[H]AMPA receptor binding in the rat hippocampal formation. Neurobiol Learn Mem 64, 257–264.

Cheng, L., Locke, C., and Davis, G.W. (2011). S6 kinase localizes to the presynaptic active zone and functions with PDK1 to control synapse development. J Cell Biol 194, 921–935.

Chiu, S.L., Diering, G.H., Ye, B., Takamiya, K., Chen, C.M., Jiang, Y., Niranjan, T., Schwartz, C.E., Wang, T., and Huganir, R.L. (2017). GRASP1 Regulates Synaptic Plasticity and Learning through Endosomal Recycling of AMPA Receptors. Neuron 93, 1405–1419 e1408.

Coba, M.P. (2019). Regulatory mechanisms in postsynaptic phosphorylation networks. Curr Opin Struct Biol 54, 86–94.

Costa-Mattioli, M., Sossin, W.S., Klann, E., and Sonenberg, N. (2009). Translational control of long-lasting synaptic plasticity and memory. Neuron 61, 10–26.

Coultrap, S.J., and Bayer, K.U. (2012). CaMKII regulation in information processing and storage. Trends Neurosci 35, 607–618.

Deutsch, E.W., Bandeira, N., Sharma, V., Perez-Riverol, Y., Carver, J.J., Kundu, D.J., Garcia-Seisdedos, D., Jarnuczak, A.F., Hewapathirana, S., Pullman, B.S., et al. (2020). The ProteomeXchange consortium in 2020: enabling ’big data’ approaches in proteomics. Nucleic Acids Res 48, D1145–D1152.

Diering, G.H., Heo, S., Hussain, N.K., Liu, B., and Huganir, R.L. (2016). Extensive phosphorylation of AMPA receptors in neurons. Proc Natl Acad Sci U S A 113, E4920–4927.

Diering, G.H., and Huganir, R.L. (2018). The AMPA Receptor Code of Synaptic Plasticity. Neuron 100, 314–329.

Diering, G.H., Nirujogi, R.S., Roth, R.H., Worley, P.F., Pandey, A., and Huganir, R.L. (2017). Homer1a drives homeostatic scaling-down of excitatory synapses during sleep. Science 355, 511–515.

Dieterich, D.C., and Kreutz, M.R. (2016). Proteomics of the Synapse--A Quantitative Approach to Neuronal Plasticity. Mol Cell Proteomics 15, 368–381.

Engholm-Keller, K., and Larsen, M.R. (2016). Improving the Phosphoproteome Coverage for Limited Sample Amounts Using TiO2-SIMAC-HILIC (TiSH) Phosphopeptide Enrichment and Fractionation. Methods Mol Biol 1355, 161–177.

Federman, N., de la Fuente, V., Zalcman, G., Corbi, N., Onori, A., Passananti, C., and Romano, A. (2013). Nuclear factor kappaB-dependent histone acetylation is specifically involved in persistent forms of memory. J Neurosci 33, 7603–7614.

Fingleton, E., Li, Y., and Roche, K.W. (2021). Advances in Proteomics Allow Insights Into Neuronal Proteomes. Front Mol Neurosci 14, 647451.

Fioravante, D., and Byrne, J.H. (2011). Protein degradation and memory formation. Brain Res Bull 85, 14–20.

Giese, K.P., and Mizuno, K. (2013). The roles of protein kinases in learning and memory. Learn Mem 20, 540–552.

Ho, V.M., Lee, J.A., and Martin, K.C. (2011). The cell biology of synaptic plasticity. Science 334, 623–628.

Hong, I., Kang, T., Yun, K.N., Yoo, Y., Park, S., Kim, J., An, B., Song, S., Lee, S., Kim, J., et al. (2013). Quantitative proteomics of auditory fear conditioning. Biochem Biophys Res Commun 434, 87–94.

Hosp, F., and Mann, M. (2017). A Primer on Concepts and Applications of Proteomics in Neuroscience. Neuron 96, 558–571.

Huganir, R.L., and Nicoll, R.A. (2013). AMPARs and synaptic plasticity: the last 25 years. Neuron 80, 704–717.

Izquierdo, I., Bevilaqua, L.R., Rossato, J.I., Bonini, J.S., Medina, J.H., and Cammarota, M. (2006). Different molecular cascades in different sites of the brain control memory consolidation. Trends Neurosci 29, 496–505.

Izquierdo, I., and Medina, J.H. (1997). Memory formation: the sequence of biochemical events in the hippocampus and its connection to activity in other brain structures. Neurobiol Learn Mem 68, 285–316.

Izquierdo, I., Quillfeldt, J.A., Zanatta, M.S., Quevedo, J., Schaeffer, E., Schmitz, P.K., and Medina, J.H. (1997). Sequential role of hippocampus and amygdala, entorhinal cortex and parietal cortex in formation and retrieval of memory for inhibitory avoidance in rats. Eur J Neurosci 9, 786–793.

Jassal, B., Matthews, L., Viteri, G., Gong, C., Lorente, P., Fabregat, A., Sidiropoulos, K., Cook, J., Gillespie, M., Haw, R., et al. (2020). The reactome pathway knowledgebase. Nucleic Acids Res 48, D498–D503.

Kahne, T., Richter, S., Kolodziej, A., Smalla, K.H., Pielot, R., Engler, A., Ohl, F.W., Dieterich, D.C., Seidenbecher, C., Tischmeyer, W., et al. (2016). Proteome rearrangements after auditory learning: high-resolution profiling of synapse-enriched protein fractions from mouse brain. J Neurochem 138, 124–138.

Kang, T., Boland, B.B., Alarcon, C., Grimsby, J.S., Rhodes, C.J., and Larsen, M.R. (2019). Proteomic Analysis of Restored Insulin Production and Trafficking in Obese Diabetic Mouse Pancreatic Islets Following Euglycemia. J Proteome Res 18, 3245–3258.

Kang, T., Boland, B.B., Jensen, P., Alarcon, C., Nawrocki, A., Grimsby, J.S., Rhodes, C.J., and Larsen, M.R. (2020). Characterization of Signaling Pathways Associated with Pancreatic beta-cell Adaptive Flexibility in Compensation of Obesity-linked Diabetes in db/db Mice. Mol Cell Proteomics 19, 971–993.

Kang, T., Jensen, P., Huang, H., Lund Christensen, G., Billestrup, N., and Larsen, M.R. (2018). Characterization of the Molecular Mechanisms Underlying Glucose Stimulated Insulin Secretion from Isolated Pancreatic beta-cells Using Post-translational Modification Specific Proteomics (PTMomics). Mol Cell Proteomics 17, 95–110.

Kempermann, G. (2019). Environmental enrichment, new neurons and the neurobiology of individuality. Nat Rev Neurosci 20, 235–245.

Kempf, S.J., Metaxas, A., Ibanez-Vea, M., Darvesh, S., Finsen, B., and Larsen, M.R. (2016). An integrated proteomics approach shows synaptic plasticity changes in an APP/PS1 Alzheimer’s mouse model. Oncotarget 7, 33627–33648.

Kim, S., and Pevzner, P.A. (2014). MS-GF+ makes progress towards a universal database search tool for proteomics. Nat Commun 5, 5277.

Kitchen, R.R., Rozowsky, J.S., Gerstein, M.B., and Nairn, A.C. (2014). Decoding neuroproteomics: integrating the genome, translatome and functional anatomy. Nat Neurosci 17, 1491–1499.

Lee, H.K. (2006). Synaptic plasticity and phosphorylation. Pharmacol Ther 112, 810–832.

Lee, H.K., Barbarosie, M., Kameyama, K., Bear, M.F., and Huganir, R.L. (2000). Regulation of distinct AMPA receptor phosphorylation sites during bidirectional synaptic plasticity. Nature 405, 955–959.

Lee, H.K., Kameyama, K., Huganir, R.L., and Bear, M.F. (1998). NMDA induces long-term synaptic depression and dephosphorylation of the GluR1 subunit of AMPA receptors in hippocampus. Neuron 21, 1151–1162.

Letunic, I., and Bork, P. (2016). Interactive tree of life (iTOL) v3: an online tool for the display and annotation of phylogenetic and other trees. Nucleic Acids Research 44, W242–W245.

Li, J., Zhang, W., Yang, H., Howrigan, D.P., Wilkinson, B., Souaiaia, T., Evgrafov, O.V., Genovese, G., Clementel, V.A., Tudor, J.C., et al. (2017). Spatiotemporal profile of postsynaptic interactomes integrates components of complex brain disorders. Nat Neurosci 20, 1150–1161.

Lisman, J., Schulman, H., and Cline, H. (2002). The molecular basis of CaMKII function in synaptic and behavioural memory. Nat Rev Neurosci 3, 175–190.

Lisman, J., Yasuda, R., and Raghavachari, S. (2012). Mechanisms of CaMKII action in long-term potentiation. Nat Rev Neurosci 13, 169–182.

Liu, H.H., McClatchy, D.B., Schiapparelli, L., Shen, W., Yates, J.R., 3rd, and Cline, H.T. (2018). Role of the visual experience-dependent nascent proteome in neuronal plasticity. Elife 7.

Lopez-Salon, M., Alonso, M., Vianna, M.R., Viola, H., Mello e Souza, T., Izquierdo, I., Pasquini, J.M., and Medina, J.H. (2001). The ubiquitin-proteasome cascade is required for mammalian long-term memory formation. Eur J Neurosci 14, 1820–1826.

Lussier, M.P., Sanz-Clemente, A., and Roche, K.W. (2015). Dynamic Regulation of N-Methyl-d-aspartate (NMDA) and alpha-Amino-3-hydroxy-5-methyl-4-isoxazolepropionic Acid (AMPA) Receptors by Posttranslational Modifications. J Biol Chem 290, 28596–28603.

Mansuy, I.M., and Shenolikar, S. (2006). Protein serine/threonine phosphatases in neuronal plasticity and disorders of learning and memory. Trends Neurosci 29, 679–686.

McNair, K., Davies, C.H., and Cobb, S.R. (2006). Plasticity-related regulation of the hippocampal proteome. Eur J Neurosci 23, 575–580.

Mi, H., Huang, X., Muruganujan, A., Tang, H., Mills, C., Kang, D., and Thomas, P.D. (2017). PANTHER version 11: expanded annotation data from Gene Ontology and Reactome pathways, and data analysis tool enhancements. Nucleic Acids Res 45, D183–D189.

Mi, H., Muruganujan, A., Casagrande, J.T., and Thomas, P.D. (2013). Large-scale gene function analysis with the PANTHER classification system. Nat Protoc 8, 1551–1566.

Milekic, M.H., Pollonini, G., and Alberini, C.M. (2007). Temporal requirement of C/EBPbeta in the amygdala following reactivation but not acquisition of inhibitory avoidance. Learn Mem 14, 504–511.

Mitsushima, D., Ishihara, K., Sano, A., Kessels, H.W., and Takahashi, T. (2011). Contextual learning requires synaptic AMPA receptor delivery in the hippocampus. Proc Natl Acad Sci U S A 108, 12503–12508.

Monroe, M.E., Shaw, J.L., Daly, D.S., Adkins, J.N., and Smith, R.D. (2008a). MASIC: A software program for fast quantitation and flexible visualization of chromatographic profiles from detected LC-MS(/MS) features. Computational Biology and Chemistry 32, 215–217.

Monroe, M.E., Shaw, J.L., Daly, D.S., Adkins, J.N., and Smith, R.D. (2008b). MASIC: a software program for fast quantitation and flexible visualization of chromatographic profiles from detected LC-MS(/MS) features. Comput Biol Chem 32, 215–217.

Montarolo, P.G., Goelet, P., Castellucci, V.F., Morgan, J., Kandel, E.R., and Schacher, S. (1986). A critical period for macromolecular synthesis in long-term heterosynaptic facilitation in Aplysia. Science 234, 1249–1254.

Murakoshi, H., Shin, M.E., Parra-Bueno, P., Szatmari, E.M., Shibata, A.C.E., and Yasuda, R. (2017). Kinetics of Endogenous CaMKII Required for Synaptic Plasticity Revealed by Optogenetic Kinase Inhibitor. Neuron 94, 37–47 e35.

Nicoll, R.A. (2017). A Brief History of Long-Term Potentiation. Neuron 93, 281–290.

Nithianantharajah, J., and Hannan, A.J. (2006). Enriched environments, experience-dependent plasticity and disorders of the nervous system. Nat Rev Neurosci 7, 697–709.

Orchard, S., Ammari, M., Aranda, B., Breuza, L., Briganti, L., Broackes-Carter, F., Campbell, N.H., Chavali, G., Chen, C., del-Toro, N., et al. (2014). The MIntAct project--IntAct as a common curation platform for 11 molecular interaction databases. Nucleic Acids Res 42, D358–363.

Palmisano, G., Parker, B.L., Engholm-Keller, K., Lendal, S.E., Kulej, K., Schulz, M., Schwammle, V., Graham, M.E., Saxtorph, H., Cordwell, S.J., et al. (2012). A novel method for the simultaneous enrichment, identification, and quantification of phosphopeptides and sialylated glycopeptides applied to a temporal profile of mouse brain development. Mol Cell Proteomics 11, 1191–1202.

Park, P., Volianskis, A., Sanderson, T.M., Bortolotto, Z.A., Jane, D.E., Zhuo, M., Kaang, B.K., and Collingridge, G.L. (2014). NMDA receptor-dependent long-term potentiation comprises a family of temporally overlapping forms of synaptic plasticity that are induced by different patterns of stimulation. Philos Trans R Soc Lond B Biol Sci 369, 20130131.

Peineau, S., Bradley, C., Taghibiglou, C., Doherty, A., Bortolotto, Z.A., Wang, Y.T., and Collingridge, G.L. (2008). The role of GSK-3 in synaptic plasticity. Br J Pharmacol 153 *Suppl 1*, S428–437.

Perez-Riverol, Y., Csordas, A., Bai, J., Bernal-Llinares, M., Hewapathirana, S., Kundu, D.J., Inuganti, A., Griss, J., Mayer, G., Eisenacher, M., et al. (2019). The PRIDE database and related tools and resources in 2019: improving support for quantification data. Nucleic Acids Res 47, D442–D450.

Qiao, H., Foote, M., Graham, K., Wu, Y., and Zhou, Y. (2014). 14-3-3 proteins are required for hippocampal long-term potentiation and associative learning and memory. J Neurosci 34, 4801–4808.

Rampon, C., Jiang, C.H., Dong, H., Tang, Y.P., Lockhart, D.J., Schultz, P.G., Tsien, J.Z., and Hu, Y. (2000). Effects of environmental enrichment on gene expression in the brain. Proc Natl Acad Sci U S A 97, 12880–12884.

Roth, R.H., Zhang, Y., and Huganir, R.L. (2017). Dynamic imaging of AMPA receptor trafficking in vitro and in vivo. Curr Opin Neurobiol 45, 51–58.

Saito, N., and Shirai, Y. (2002). Protein kinase C gamma (PKC gamma): function of neuron specific isotype. J Biochem 132, 683–687.

Schanzenbacher, C.T., Langer, J.D., and Schuman, E.M. (2018). Time- and polarity-dependent proteomic changes associated with homeostatic scaling at central synapses. Elife 7.

Schwammle, V., and Jensen, O.N. (2010). A simple and fast method to determine the parameters for fuzzy c-means cluster analysis. Bioinformatics 26, 2841–2848.

Shipton, O.A., and Paulsen, O. (2014). GluN2A and GluN2B subunit-containing NMDA receptors in hippocampal plasticity. Philos Trans R Soc Lond B Biol Sci 369, 20130163.

Spiegel, I., Mardinly, A.R., Gabel, H.W., Bazinet, J.E., Couch, C.H., Tzeng, C.P., Harmin, D.A., and Greenberg, M.E. (2014). Npas4 regulates excitatory-inhibitory balance within neural circuits through cell-type-specific gene programs. Cell 157, 1216–1229.

Szklarczyk, D., Morris, J.H., Cook, H., Kuhn, M., Wyder, S., Simonovic, M., Santos, A., Doncheva, N.T., Roth, A., Bork, P., et al. (2017). The STRING database in 2017: quality-controlled protein-protein association networks, made broadly accessible. Nucleic Acids Research 45, D362–D368.

Tadi, M., Allaman, I., Lengacher, S., Grenningloh, G., and Magistretti, P.J. (2015). Learning-Induced Gene Expression in the Hippocampus Reveals a Role of Neuron -Astrocyte Metabolic Coupling in Long Term Memory. PLoS One 10, e0141568.

Taubenfeld, S.M., Milekic, M.H., Monti, B., and Alberini, C.M. (2001). The consolidation of new but not reactivated memory requires hippocampal C/EBPbeta. Nat Neurosci 4, 813–818.

Taus, T., Kocher, T., Pichler, P., Paschke, C., Schmidt, A., Henrich, C., and Mechtler, K. (2011). Universal and Confident Phosphorylation Site Localization Using phosphoRS. Journal of Proteome Research 10, 5354–5362.

Thingholm, T.E., Jorgensen, T.J., Jensen, O.N., and Larsen, M.R. (2006). Highly selective enrichment of phosphorylated peptides using titanium dioxide. Nat Protoc 1, 1929–1935.

Thingholm, T.E., and Larsen, M.R. (2009). The use of titanium dioxide micro-columns to selectively isolate phosphopeptides from proteolytic digests. Methods Mol Biol 527, 57–66, xi.

Thomas, G.M., and Huganir, R.L. (2004). MAPK cascade signalling and synaptic plasticity. Nat Rev Neurosci 5, 173–183.

Thygesen, C., Boll, I., Finsen, B., Modzel, M., and Larsen, M.R. (2018). Characterizing disease-associated changes in post-translational modifications by mass spectrometry. Expert Rev Proteomics 15, 245–258.

Thygesen, C., Larsen, M.R., and Finsen, B. (2019). Proteomic signatures of neuroinflammation in Alzheimer’s disease, multiple sclerosis and ischemic stroke. Expert Rev Proteomics 16, 601–611.

Tully, T., Preat, T., Boynton, S.C., and Del Vecchio, M. (1994). Genetic dissection of consolidated memory in Drosophila. Cell 79, 35–47.

Vianna, M.R., Szapiro, G., McGaugh, J.L., Medina, J.H., and Izquierdo, I. (2001). Retrieval of memory for fear-motivated training initiates extinction requiring protein synthesis in the rat hippocampus. Proc Natl Acad Sci U S A 98, 12251–12254.

Villareal, G., Li, Q., Cai, D., and Glanzman, D.L. (2007). The role of rapid, local, postsynaptic protein synthesis in learning-related synaptic facilitation in aplysia. Curr Biol 17, 2073–2080.

Whitlock, J.R., Heynen, A.J., Shuler, M.G., and Bear, M.F. (2006). Learning induces long-term potentiation in the hippocampus. Science 313, 1093–1097.

Woolfrey, K.M., and Dell’Acqua, M.L. (2015). Coordination of Protein Phosphorylation and Dephosphorylation in Synaptic Plasticity. J Biol Chem 290, 28604–28612.

Xu, Y., Song, X., Wang, D., Wang, Y., Li, P., and Li, J. (2021). Proteomic insights into synaptic signaling in the brain: the past, present and future. Mol Brain 14, 37.

Zalcman, G., Federman, N., Fiszbein, A., de la Fuente, V., Ameneiro, L., Schor, I., and Romano, A. (2018). Sustained CaMKII Delta Gene Expression Is Specifically Required for Long-Lasting Memories in Mice. Mol Neurobiol.

Zhang, Y., Cudmore, R.H., Lin, D.T., Linden, D.J., and Huganir, R.L. (2015). Visualization of NMDA receptor-dependent AMPA receptor synaptic plasticity in vivo. Nat Neurosci 18, 402–407.

Zhang, Y., Fukushima, H., and Kida, S. (2011). Induction and requirement of gene expression in the anterior cingulate cortex and medial prefrontal cortex for the consolidation of inhibitory avoidance memory. Mol Brain 4, 4.

